# PRO-FitS: a novel phenotypic assay to identify enhancers of proteostasis in *C. elegans*

**DOI:** 10.64898/2026.01.22.701023

**Authors:** Daniela Vilasboas-Campos, Jorge Humberto Fernandes, Marta Daniela Costa, Joana Pereira-Sousa, Joana Lopes, Liliana Costa-Meireles, Bruna Ferreira-Lomba, Jorge Diogo Da Silva, Patrícia Maciel, Andreia Teixeira-Castro

**Author notes:** corresponding authors Requests for further information and resources should be directed to Andreia Teixeira-Castro.

## Abstract

The prevalence of neurodegenerative diseases (NDs) continues to rise with the aging of populations worldwide, representing a pressing need for the establishment of therapeutic strategies. Maintaining proteostasis is crucial for healthy aging, as the accumulation of misfolded and aggregated proteins is a key contributor to age-related cellular dysfunction and disease. This study introduces a novel phenotypic assay using *Caenorhabditis elegans* to screen for small molecule enhancers of proteostasis, aiming at mitigating the proteotoxic stress associated with NDs. This new methodology- PRO-FitS- uses *C. elegans* motor activity as a proxy for the PROteome Fitness State upon a noxious protein-denaturating stimulus, while allowing a fast and experimenter-free readout. We demonstrate the efficacy of the assay by validating the role of pharmacological mTOR inhibition and serotonergic signaling activation in reducing heat shock-induced proteotoxic damage at the whole-organism level. PRO-FitS will allow the identification of novel compounds that alleviate protein aggregation disorders, potentially revealing new pathways and cellular targets not previously implicated in proteotoxicity.

**Significance Statement:** Neurodegenerative diseases remain without effective cures, in part due to the lack of scalable methods to identify compounds that improve proteostasis. We developed PRO-FitS, a whole-organism, automated phenotypic assay in *C. elegans* that uses motor activity recovery as a proxy for proteome fitness after proteotoxic stress. This platform enables rapid, unbiased screening of small molecules and genetic modifiers, bridging the gap between cellular assays and complex animal models. By demonstrating the assay’s robustness in both wild-type and disease-relevant contexts, we establish PRO-FitS as a versatile tool for discovering therapeutic candidates and uncovering novel pathways relevant to protein aggregation disorders.

## Introduction

Neurodegenerative diseases (NDs) including Alzheimer’s, Parkinson’s, Huntington’s and other polyglutamine diseases, represent a significant challenge in the field of medical science, primarily due to their progressive and irreversible impact on brain and organism function. These disorders are characterized by the deleterious accumulation of misfolded proteins, leading to proteotoxic stress, cellular dysfunction, and death (*1*, *2*). As the global population ages, the prevalence of these diseases increases, underscoring the urgent need to identify novel therapeutic approaches - currently lacking - to stall or prevent disease progression (*3*, *4*).

Key factors for the onset and progression of NDs are pathological protein aggregation and the disruption of protein homeostasis, or proteostasis (*5*). Contributors to proteostasis imbalance include genetic mutations, environmental factors (or exposome), the age-related decline in proteostasis maintenance, competition for limited proteostasis resources, dysregulated protein degradation pathways, and defects in cellular non-autonomous signaling or inter-tissue communication (*6–11*). These interconnected processes drive the accumulation of toxic protein aggregates - characteristic of NDs.

Proteostasis is achieved when proteins are correctly synthesized, folded, trafficked, and timely degraded, essential mechanisms for maintaining the proteome in a healthy state (*12*, *13*). The cellular components that contribute to maintain proteostasis enclose the proteostasis network. Currently, nine main divisions of the proteostasis network are considered, comprising the mitochondrial, endoplasmic reticulum, cytonuclear and nuclear proteostasis, cytosolic translation, proteostasis regulation, extracellular proteostasis, the ubiquitin-proteasome system, and the autophagy-lysosome pathway, collectively comprising approximately 3000 players (*14*, *15*). A growing body of evidence from invertebrate model systems suggests that proteostasis is coordinated systemically across different cells and tissues. To counteract proteotoxic stress, cells activate highly conserved protective mechanisms, including the heat shock response (HSR) and unfolded protein responses (UPRs) specific to different cellular compartments. These responses attempt to monitor the cellular environment and restore proteostasis, alleviating the burden of misfolded proteins (*16–19*). In *C. elegans,* the HSR is regulated by thermosensory neurons that sense thermal stress and activate heat shock factor 1 (HSF-1)-dependent transcriptional programs, elevating the levels of molecular chaperones to prevent misfolding and guide abnormal proteins to degradation (*20*, *21*). Additional studies have highlighted the crucial role of the nervous system in coordinating not only the HSR but also mitochondrial and endoplasmic reticulum UPRs in peripheral tissues, particularly in the gut, through the release of neuropeptides and neurotransmitters (*21–27*). Tissues experiencing proteotoxic stress also induce transcellular chaperone signaling that activates molecular chaperones, such as hsp-90, in distal tissues (*28*, *29*). This evidence indicates that the proteome status in one tissue can be communicated to other tissues, providing whole-organism protection and contributing to survival and longevity (*10*, *30*).

Despite significant efforts, the identification of effective therapies targeting the proteotoxicity inherent to NDs remains a challenge. The development of novel and improved screening methods to identify drugs that enhance proteostasis - by reducing time, minimizing experimenter dependency, and providing easily quantifiable and consistent readouts - holds promise for uncovering new therapeutic options for NDs with a proteotoxic component.

*C. elegans* is a valuable model to study several NDs and conduct drug screenings, benefiting from its biological complexity as a multicellular organism, conservation of core biochemical features, a well-defined neuronal connectome, representativeness of the majority of the neurotransmitter systems, and associated complex behaviors (*31–34*). This model offers a unique opportunity to systematically explore and understand proteostasis mechanisms, providing an ideal platform to manipulate its complex responses to environmental and endogenous stressors, such as heat shock (HS), and thereby uncover potential treatments for NDs characterized by disruptions in proteostasis (*32*, *35–38*).

Here, we describe a novel phenotypic assay to screen for enhancers of proteostasis in *C. elegans*. In this approach, acute thermal stress challenges the organism’s proteome, affecting animals’ motor activity in a robust, objective, and quantifiable manner. As a proof-of-concept in wild-type (WT) animals, we demonstrate that known pharmacological and genetic modulators of proteostasis influence HS-induced proteome disruption, resulting in a faster recovery of the animals’ motor activity. We further validate the methodology by testing it in animals expressing abnormal polyglutamine or tau proteins in their nervous system, moving the assay into ND-relevant contexts. Additionally, we demonstrate the adaptability of the approach to capture distinct thermotolerance outcomes. In this way, we propose a robust methodology for identifying drugs that can mitigate protein dyshomeostasis and subsequent motor dysfunction, offering a promising platform for discovering novel therapeutic candidates to address proteinopathies.

## Results

### Development of a phenotypic assay to screen for chemical enhancers of proteostasis in *C. elegans*

*C. elegans* is a useful whole-organism platform for small molecule screens to identify compounds that modify complex phenotypes, such as motor behavior (*39*, *40*). Here, we hypothesized that organismal motor behavior alterations in response to a heat shock (HS) noxious stimulus serve as a proxy to monitor changes in proteome fitness and proteostasis dynamics, allowing the identification of chemical modifiers of nervous system-relevant proteotoxic stress, with impact on NDs associated with protein aggregation (Fig. 1A). The effect of HS on animals’ motor activity in miniaturized liquid cultures can be quantified in an easy and automated manner using a computerized plate reader (*41*), in which the detection of activity is based on infrared microbeam interruptions. Animal movement across the light beams generates a transient fluctuation in the signal received by the phototransistors, and worm activity is detected by digital analysis of the phototransistor output (*41*). For these assays, synchronized animals of the reference strain N2 (egg stage- day 0) were allowed to grow for three days in the presence of food (*42*). On day 3, we measured the animals’ locomotor activity (basal activity), after which we induced HS, and monitored activity right after HS (post-HS activity) and during a recovery period (total recovery activity and final recovery activity) (Fig. 1B).

**Figure 1.**
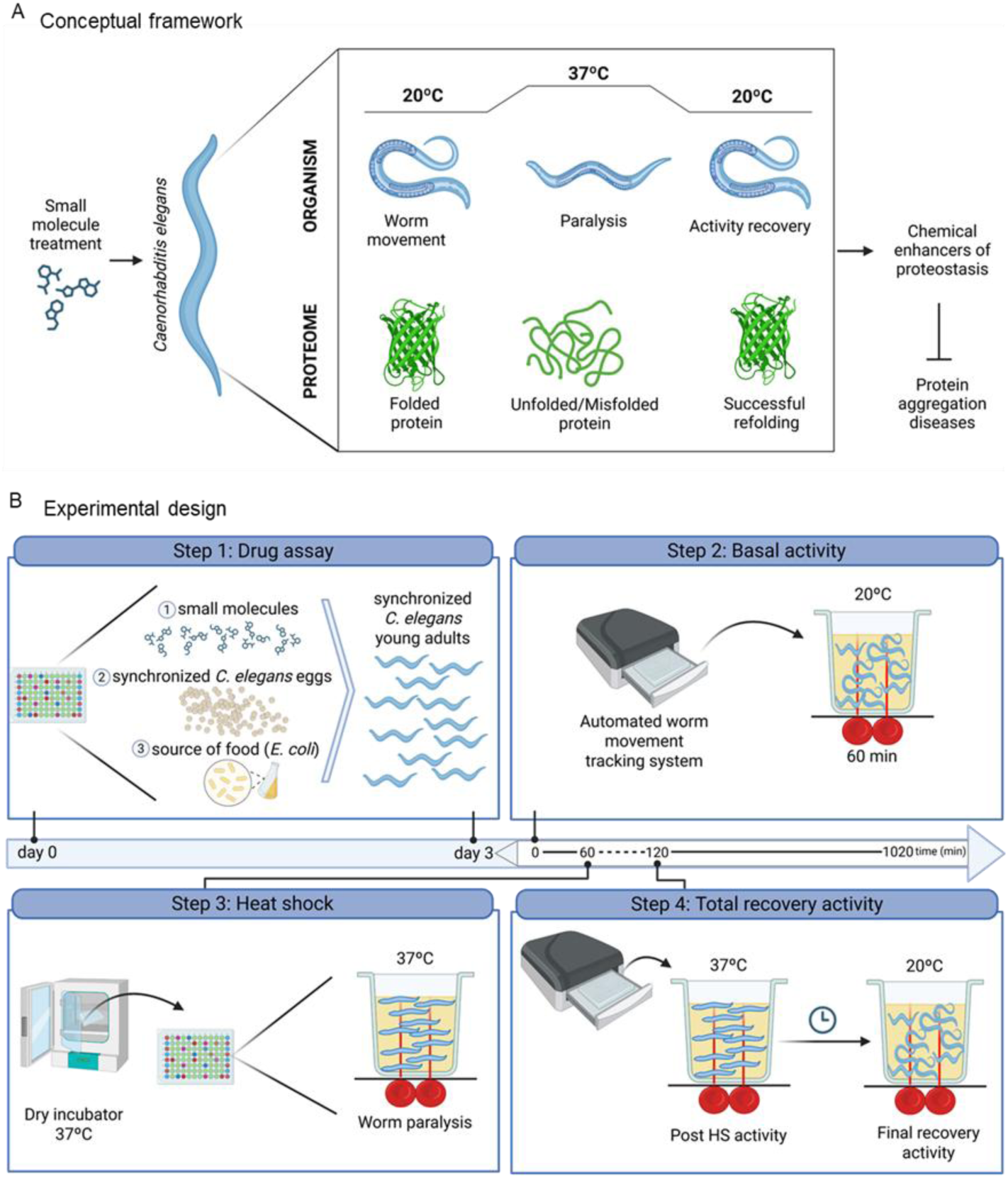
Assay framework to identify chemical enhancers of proteostasis with potential activity against protein aggregation disorders in *C. elegans*. **(A)** HS-induced alterations in animals’ motor behavior are used as a proxy of the proteome fitness state. At 20 °C, the worms move normally with properly folded proteins. When subjected to HS at 37 °C, the animals undergo paralysis likely associated with protein misfolding. After returning to 20 °C, the animals gradually recover movement, suggesting protein refolding. This approach enables the rapid and experimenter-free identification of potential therapeutic agents for protein aggregation disorders. **(B)** Experimental pipeline development to identify chemical modulators of proteostasis in *C. elegans*. The methodology involves four critical steps: 1) Performing a drug assay to expose synchronized eggs of WT *C. elegans* to small molecules, at a range of concentrations (day 0); 2) Assessing basal activity using an automated worm movement tracking system to measure pre-HS locomotor activity of young adult worms (day 3 post-hatching) at 20 °C for 1 hour; 3) Proteotoxic stress induction by placing the plates in a 37 °C incubator for a time period (e.g. 60 min); and 4) Monitoring post-HS recovery activity to determine the efficacy of the compounds in restoring normal locomotory activity of the animals, after a noxious stimulus.

During assay optimization, we evaluated different HS durations from 30 to 120 min at 37 °C (Fig. 2). We aimed to identify an HS exposure condition that disrupted motor function without causing irreversible damage, allowing animals to recover over time. This balance was essential to develop a dynamic assay for monitoring proteostasis resilience and recovery. As expected, longer HS durations were associated with a more severe motor activity phenotype immediately following HS and during recovery phases (Fig. 2A). After 30 or 45 min of HS, animals’ locomotor activity was significantly reduced by more than 50% compared to basal activity levels, whereas HS of 60 min or more caused animals’ paralysis (Fig. 2, A and B). With 60 min of HS, the animals recovered nearly baseline levels within 720 to 1020 min (corresponding to 10 to 15h post-HS). In contrast, after 90 or 120 min of HS, animals showed limited motor recovery (23 and 7% of recovery, respectively), as seen by the comparison of the final recovery activity, which averages the activity of the animals in the last four time-points of recovery (Fig. 2C). A distinct recovery profile among the different HS duration protocols was also seen when comparing the cumulative activity of the animals across the entire recovery period (total recovery activity) (Fig. 2D). All data measures were shown as a ratio to basal activity (Fig. 2, B-D). This ensured that any potential differences in motor behavior after HS were not due to pre-existing changes in activity, potentially not related to proteotoxic stress. Although significant variability among independent assays was registered, as extensively described previously in the literature in other HS-based assays (*43*, *44*), the dynamic range of the recovery from a 60 min exposure to HS can allow the identification of proteostasis modulators, by measuring the time required to achieve 50% recovery of the basal activity, in the presence or absence of a given compound, which we called T50 (Fig. 2E).

**Figure 2.**
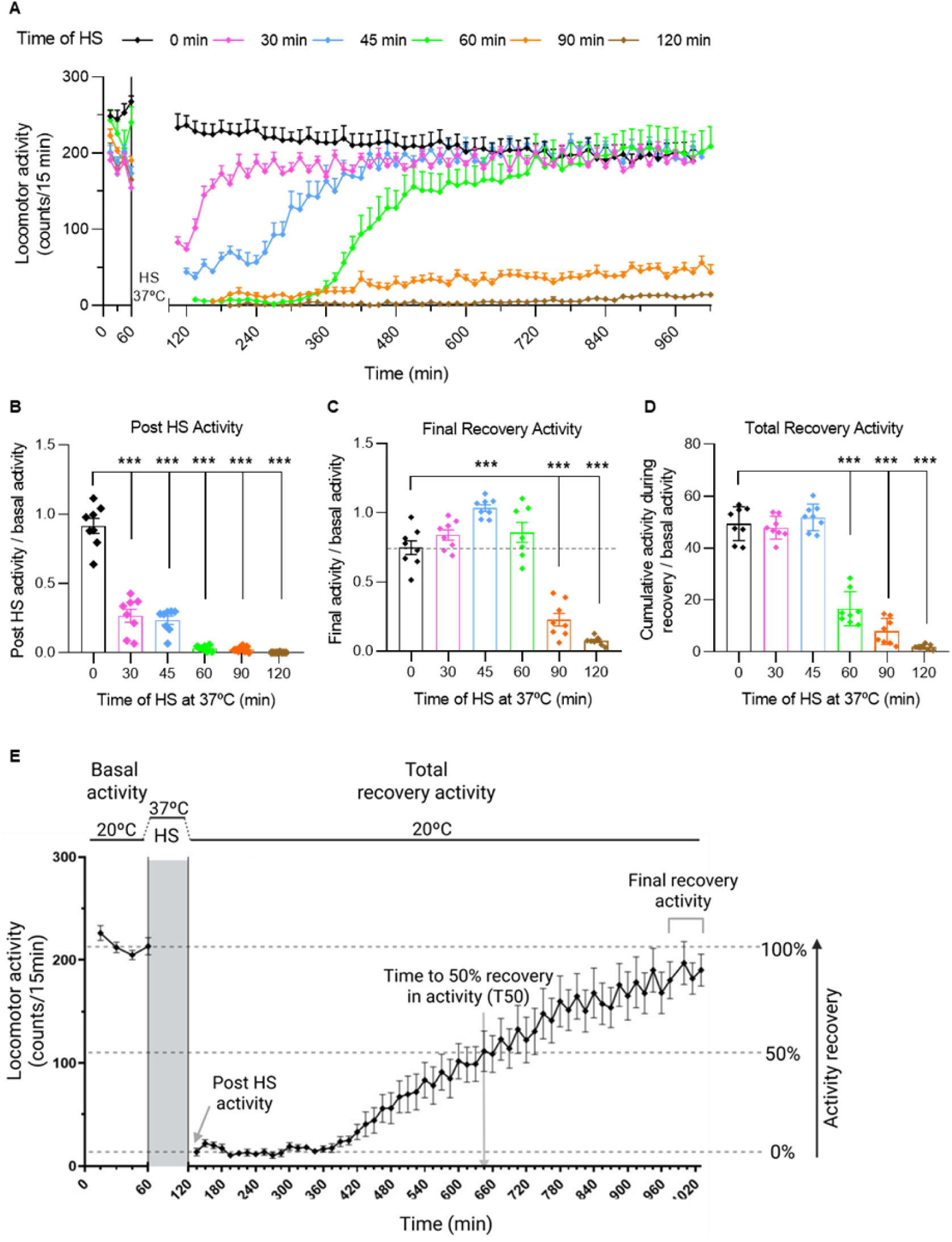
Effect of increasing HS durations on locomotor activity of WT C*. elegans.* **(A)** Locomotor activity of WT *C. elegans* under normal growth conditions (0 min of HS) and following 30, 45, 60, 90, and 120 min of HS at 37 °C, measured using an automated worm movement tracking system. **(B)** Post-HS activity 15 min after the removal of the stressor and normalized to mean basal activity levels, demonstrating that only HS durations of 60 min or longer resulted in complete paralysis of the worms. **(C)** Final recovery activity of the worms, normalized to mean basal activity, indicating that HS durations of up to 60 min allow animals to restore their motor activity to near baseline levels. Final recovery activity was determined by averaging the motor activity values from the last four timepoints of analysis. **(D)** Total recovery activity was determined by calculating the total cumulative activity of the animals measured over a 15-hour recovery period post-HS, normalized to mean basal activity. Graphs show the mean ± standard deviation (SD) of locomotor activity measures across eight wells per condition (N=400 animals) and are representative of at least three independent experiments. P values (P < 0.001 (***)) were calculated using One-way ANOVA, Tukey’s multiple comparisons test. **(E)** Graphical representation of the phenotypic assay to identify chemical enhancers of proteostasis, illustrating the basal activity, 1h HS exposure at 37 °C, and the subsequent recovery phase of motor activity: post-HS activity, final recovery activity and total recovery activity. The time taken to recover 50% of the basal activity upon HS (T50) serves as a measure to identify chemical compounds with proteostasis-enhancing activity.

A key aspect of assay development is the robustness of the measures used, namely their resistance to expectable experimental error. Quality control tests confirmed that differences of 10 animals among wells, within the range of 40-60 animals per well, did not significantly alter activity counts (fig. S1, A and B), and bacterial optical density (OD) values higher than 0.3 were established as optimal to ensure normal animal growth and movement without affecting experimental outcomes (fig. S1, C and D). Moreover, their activity measurements remained consistent across wells (under the same conditions), regardless of the position on the plate (fig. S1, E to H). Importantly, despite the lack of agitation, no hypoxia was induced throughout the motility assessments (fig. S2). Indeed, *hif-1* mutant animals, which show increased susceptibility to low oxygen levels (*45*), remained active and healthy for 24 hours in the plate reader (fig. S2A). Stable pH levels were observed when animals were kept in the plate reader with or without agitation (normal growth conditions inside the shaker or during activity measurements, respectively) (fig. S2B). Furthermore, the absence of activation in hypoxia-induced HIF-1-dependent transcriptional reporter strain (*nhr-57*p::GFP) (fig. S2, C and D) also indicated no adverse effects from the 14-hour activity recordings during the post-HS recovery phase. Collectively, these findings ensure the assay’s reliability for extended post-HS activity recordings and exclude differences in resistance to hypoxia as a potential confounder.

Next, we asked whether the impact of HS on locomotor activity (Fig. 2E) was associated with the predicted transient increase in the animals’ protein aggregation load (Fig. 3). Whole-organism filter retardation assay showed an increase in amyloid fibrils and oligomers immediately after HS, identified with an anti-amyloid fibrils’ antibody (*46*). Importantly, at four hours post-HS, there was already a significant decrease in the aggregated species detected by this antibody (Fig. 3A and fig. S3). To further test if 1h of HS at 37 °C was sufficient to cause a recoverable proteome dyshomeostasis *in vivo*, we used the extracellular aggregation reporter strain of secreted LBP-2 (*47*), depicted in fig. S4. Confocal laser scanning imaging (fig. S4, A and B) quantified LBP-2::tagRFP protein foci and replicated previous findings on the impact of animals’ age on extracellular protein aggregation (fig. S4, C and E). Temporal analysis of aggregation patterns suggested day 6 as a suitable timepoint for further studies, as most animals showed mild aggregation profiles, susceptible to being further challenged by a thermal insult (fig. S4, C-E). Following HS, aggregation of LBP-2::tagRFP in the anterior bulb of the animals significantly increased and returned to near basal levels 4 to 12 hours post-HS (Fig. 3, B and C). At 12h post-HS, the aggregation profile resembled that of non-HS animals of the same age (Fig. 3B, 12h-NoHS). Taken together, these findings suggest that the locomotor activity of WT *C. elegans* post-heat stress and during recovery can be used as a readout for whole-organism proteome fitness. Therefore, this assay was named PRO-FitS for its ability to evaluate the proteome fitness state.

**Figure 3.**
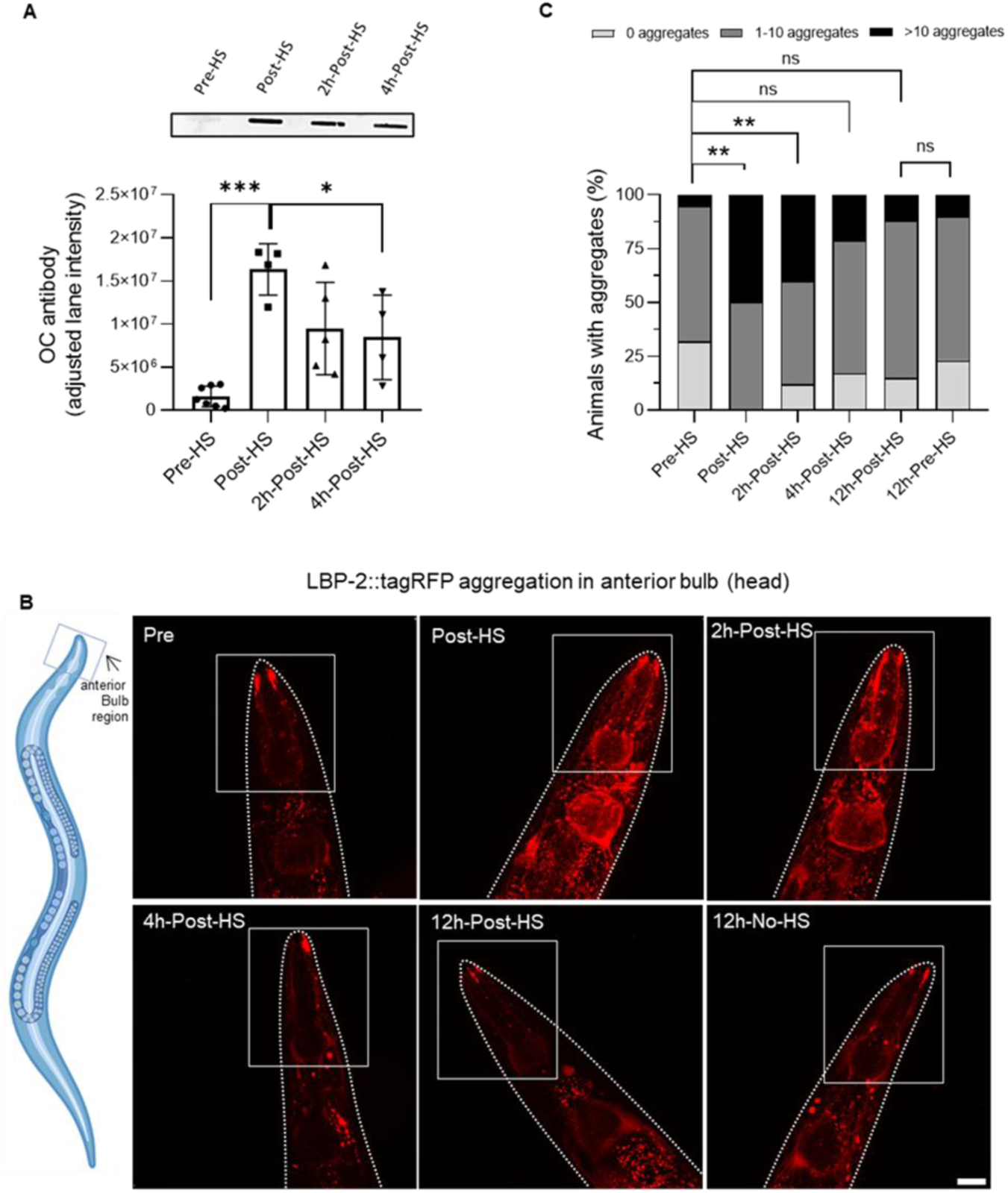
Impact of HS on the *C. elegans* proteome. **(A)** Filter retardation assay showing the amount of amyloid fibrils at pre- and post-HS timepoints. Data on the graph represents the mean of amyloid fibrils signals on the membrane (Anti-Amyloid Fibrils OC Antibody), normalized to the total protein signals (AzureRed® stained gel). P values (P < 0.05) were determined by Chi-Square test. Each bar represents mean values ± SD of 4-7 biological replicates with around 2000 animals in each. **(B)** Representative fluorescence microscopic images of LBP-2::tagRFP aggregation (day 6) in the anterior bulb region (head region highlighted in the *C. elegans* schematic representation) at different timepoints of the assay. Scale bar = 20 µm. **(C)** Quantification of LBP-2::tagRFP aggregation patterns at different assay timepoints (day 6). P values (P < 0.05) were determined by Chi-Square test. Each bar represents mean values ± SD of 4-5 independent experiments with at least 10 animals per condition in each.

### Assay validation - temsirolimus (CCI-779), an inhibitor of mTOR, enhances proteostasis in WT *C. elegans* upon heat shock

Next, and as a first validation of the potential of PRO-FitS to detect chemical enhancers of proteostasis, we tested whether a known modulator of protein aggregation *in vivo* would increase the capacity of WT animals to recover their motor performance upon HS. Therefore, we evaluated the effect of temsirolimus (CCI-779), a synthetic rapamycin ester that inhibits the mammalian target of rapamycin (mTOR) (*48–51*). This molecule was previously shown to activate protein degradation through autophagy (*50*), being particularly potent upon acute proteotoxic stress (*52*).

After conducting compound safety assays to establish a range of safe concentrations of CCI-779 to be administered to WT *C. elegans* (fig. S5A), we determined the baseline of motor activity of WT animals treated with vehicle or CCI-779 (1 pM-100 µM). After 3 days of treatment, no significant differences were noted in the basal activity of the animals under standard growth conditions (Fig. 4A and fig. S5B). Post-HS activity measures demonstrated a drastic reduction in motor activity, with vehicle-treated animals showing almost complete paralysis, underscoring the severe impact of the HS protocol. This effect served as a benchmark to evaluate the protective effects of CCI-779 (Fig. 4, A to D and movie S1) and showed a significant increase in the ratio of post-HS activity to basal activity at concentrations higher than 0.1 nM (Fig. 4B), suggesting an increased resilience to HS-induced damage upon treatment. In fact, animals treated with CCI-779 showed a dose-dependent improvement in recovery post-HS. At a concentration of 100 µM, treated animals began showing signs of recovery as early as at 150-165 min. This was faster than the control group, which started to recover at 285-315 min. The rapid recovery observed in the CCI-779-treated groups highlights its potential to mitigate HS-induced damage (Fig. 4A). Over 15 hours following HS, both treated and untreated groups recovered fully from the proteotoxic stress (fig. S5C). However, CCI-779-treated animals displayed significantly greater total recovery activity, suggesting a higher overall resilience to HS-induced damage (Fig. 4C), in addition to a faster recovery initiation. The T50, representing the time required for the population to achieve 50% of the basal recovery activity, was markedly reduced in CCI-779-treated animals relative to the control group (Fig. 4D), with CCI-779 at 100 µM showing the highest effect size (fig. S5D). This indicates a more efficient recovery process upon treatment, further supporting the effectiveness of CCI-779 in enhancing proteostasis under stress conditions.

**Figure 4.**
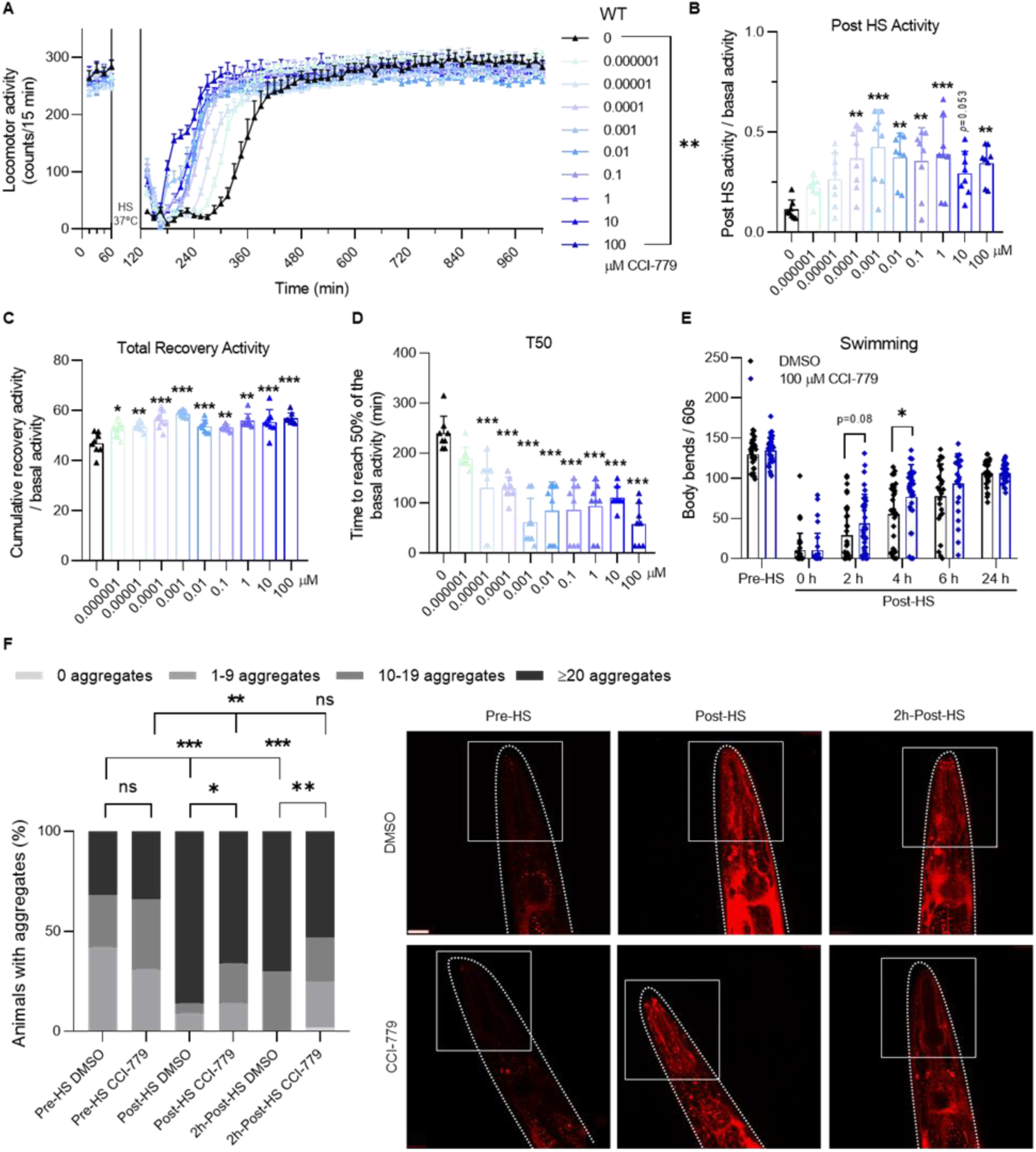
Impact of temsirolimus (CCI-779- mTOR pathway inhibitor) treatment on motor behavior and aggregation of *C. elegans* upon a noxious heat stress. **(A)** Motor activity of WT *C. elegans* treated for three days with different concentrations of CCI-779, evaluated in the WMicrotracker at pre- and post-HS and during recovery phases. Data presented as mean ± SD are representative of at least three independent trials. In each experiment, the motor activity of 400 animals per condition (8 wells/condition) was evaluated. *P* value (*P* < 0.05) was calculated using Two-way RM ANOVA, Tukey’s multiple comparisons test. **(B)** Post-HS activity- assessed for 15 min after the removal of the HS stimulus- divided by the mean of the basal activity of each experimental group. **(C)** Total recovery activity of the animals measured over a 15-hour recovery period post-HS (normalized to basal activity). **(D)** Time to achieve 50% recovery of the basal activity after HS (T50) of CCI-779- and vehicle-treated animals. ((B-D) Data on the graphs represent mean ± SD values of the activity of 400 animals per condition. Data is representative of at least three independent trials. *P* values were calculated using One-way ANOVA, Tukey’s multiple comparisons test. **(E)** Locomotor activity quantified by body bends per minute, comparing baseline activity levels (pre-HS) to levels immediately following HS and at various recovery intervals (2-, 4-, 6-, and 24-hour post-HS). A total of approximately 30 animals were assayed per condition, across 3 independent experiments. Statistical significance levels (*P* values) were calculated using One-way ANOVA, Tukey’s multiple comparisons test. (**F**) Quantification of LBP-2::tagRFP aggregation load post-HS and upon 2h of recovery (day 4). *P* values (*P* < 0.05) were determined by Chi-Square test. Each bar represents mean values ± SD of 4 independent experiments with at least 8 animals per condition in each. Scale bar = 20 µm.

Manual counting of body bends (*53*, *54*) at pre-HS, post-HS, and during recovery phases confirmed the findings of this assay through an orthogonal method, CCI-779-treated animals exhibiting a higher number of body bends 2 to 4 hours post-HS compared to the control group (Fig. 4E and movie S2). However, the automated nature of the PRO-FitS assay, allowing continuous activity assessment, proved more advantageous, offering greater sensitivity, faster execution, and minimal reliance on experimenter involvement.

Aggregation of LBP-2::tagRFP proteins in the anterior bulb of the animals significantly increased upon HS, although slightly less when treated with CCI-779 (Fig. 4F). Importantly, at 2h post-HS, CCI-779-treated animals showed a significant reduction in the percentage of animals with aggregates, to levels not different from pre-HS animals (Fig. 4F), phenocopying the activity recovery of WT animals upon CCI-779 administration (Fig. 4A).

Taken together, these results constitute a proof-of-principle that motor behavior alterations in response to HS serve as a proxy to monitor changes in proteome fitness, while allowing the identification of chemical modifiers of proteotoxic stress.

### Applicability of PRO-FitS to drug testing in a neurodegenerative disease (ND) model

Next, we explored whether this new methodology could be applied to *C. elegans* strains that express aggregation-prone proteins. The fact that treatment with CCI-779 and other rapamycin analogs alleviated pathology and associated toxicity in animal models of several NDs, including the suppression of Spinocerebellar ataxia type 3 (SCA3) or Machado-Joseph disease (MJD) phenotypes (*48–50*, *55–57*), prompted us to test CCI-779 in a well-established *C. elegans* model of MJD. This model expresses human ATXN3 with 130 glutamine residues (AT3Q130) and mimics the misfolding and aggregation of the human disease-causing protein, while challenging proteostasis and disrupting cellular function (*58*).

Chronic treatment with CCI-779, as anticipated, alleviated locomotion deficits in AT3Q130 animals during the motility test (Fig. 5A), a standard assay based on the crawling ability in solid medium, used in this model to evaluate potential therapeutic strategies (*57*, *59*). This result provides proof-of-concept evidence for assessing whether CCI-779 accelerated activity recovery in AT3Q130 animals following HS.

**Figure 5.**
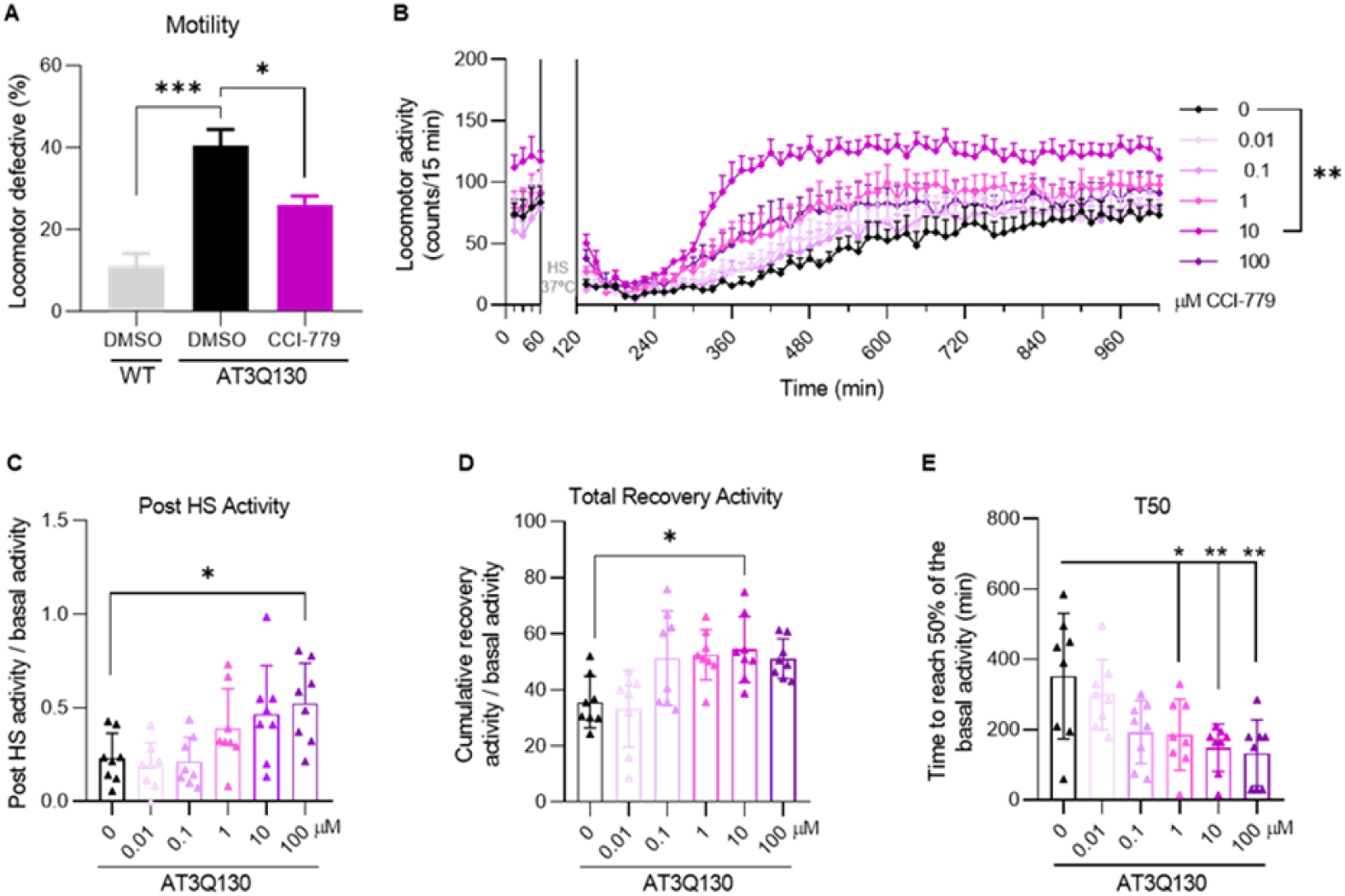
Motor recovery upon HS of CCI-779-treated *C. elegans* model of Machado-Joseph disease (MJD)/Spinocerebellar Ataxia type 3 (SCA3). **(A)** Motility analysis of AT3Q130 animals treated with CCI-779 (10 µM). **(B)** Measurement of post-HS activity in CCI-779 treated and untreated animals. Each biological experiment included eight technical replicates (N=400) for each concentration. *P* values were calculated using Two-way RM ANOVA, Tukey’s multiple comparisons test. The data are presented as mean ± SD. **(C)** Post-HS activity- assessed for 15 min after the removal of the HS stimulus- divided by the mean of the basal activity of each experimental group.**(D)** Total recovery activity of the animals measured over a 15-hour recovery period post-HS (normalized to basal activity). **(E)** Time to achieve 50% recovery of the basal activity after HS (T50) of CCI-779- and vehicle-treated animals. (C-E) Data represent mean ± SD values of the activity of 400 animals per condition. Data is representative of at least three independent trials. *P* values were calculated using One-way ANOVA, Tukey’s multiple comparisons test.

Experiments to determine the optimal HS duration for AT3Q130-expressing animals demonstrated that, similarly to WT animals, HS periods of 90 and 120 min were too severe for this strain. These longer HS durations hindered subsequent activity recovery of the animals. On the other hand, shorter HS periods of 45 and 60 min allowed for significant activity recovery, with the animals reaching baseline activity levels within 15 hours post-HS. These findings suggested that moderate HS durations are optimal for assessing proteostasis dynamics in this model, while still permitting recovery of motor function (fig. S6, A-C). Moreover, it was possible to discriminate the motor phenotype differences among WT and transgenic *C. elegans* expressing non-pathogenic (AT3Q75) and pathogenic ataxin-3 proteins (AT3Q130) when measuring the basal activity in the automated plate reader. However, immediately after an HS of 60 min, there was no significant difference in the motor response between these genotypes, and their final recovery activity was similar to their basal motor activity profiles, further suggesting that an HS of 60 min is also suitable for these animals (fig. S6, D-G).

While AT3Q130 animals treated with CCI-779 at different concentrations showed no significant changes in basal activity (fig. S6H), treatment at concentrations >1 µM elicited an earlier recovery initiation from HS-induced motor dysfunction, when compared to the control group. Specifically, the onset of recovery for treated animals occurred between 240- and 300-min post-HS, while controls exhibited the first recovery signs between 360- and 420-min (Fig. 5B). Moreover, higher concentrations of CCI-779 markedly prevented HS-related damage, reaching statistical significance at 100 µM of CCI-779 (Fig. 5C), while maintaining motor activity counts above baseline levels throughout the 15-hour recovery period at 10 µM treatment concentration (Fig. 5D). By the end of the observation period, all treated and untreated animals returned to baseline activity levels (fig. S6I). Notably, the recovery profile of CCI-779-treated animals (at 10 µM) paralleled that of the non-pathogenic AT3Q75-expressing animals, with no significant differences in curve comparison during the HS recovery period (fig. S6J).

T50 comparisons showed that AT3Q130 animals treated with 1, 10 and 100 µM of CCI-779 significantly outperformed vehicle-treated animals. The T50 for treated animals at 10 µM was 188 min, compared to 373 min in the control group, confirming the efficacy of this drug in promoting faster motor recovery upon HS of MJD animals (Fig. 5E and fig. S6K).

### Enhanced heat shock resilience of *C. elegans* with increased serotonergic signaling strengthens the utility of the assay for the identification of proteostasis modulating genes

Whole-organism inter-tissue signaling pathways are critical during conditions of stress, where dysfunction in certain organs may contribute to the loss of inter-tissue communication pathways and homeostasis (*60*, *61*). The nervous system regulates the activity of other tissues through hormones or neuronal signaling (e.g. neurotransmitters and neuropeptides), but also the state of protein homeostasis. Serotonergic signaling was widely described as being involved in this mechanism, its release from a single neuron being sufficient to suppress protein aggregation in distal tissues (*20*). Moreover, the communication of mitochondrial-mediated proteotoxic stress between neurons and intestinal cells is also mediated by serotonin (*25–27*). Based on these findings and the fact that blocking the serotonin transporter MOD-5/SERT suppressed mutant ataxin-3 protein aggregation (*62*), we hypothesized that whole-organism modulation of serotonergic signaling would result in increased resilience to HS, placing *mod-5* as a proteostasis-suppressor gene.

Treatment with the selective serotonin reuptake inhibitor citalopram (CIT), previously shown to target MOD-5/SERT (*63*), significantly improved the recovery kinetics in WT *C. elegans* post-HS, particularly at a concentration of 0.001 µM (fig. S7 and movie S3). While basal activity and final recovery activity of the animals in the PRO-FitS assay were unchanged upon CIT treatment (fig. S7B and E), post-HS and total recovery activities increased significantly (fig. S7C and D). Recovery time lengths, as measured by the T50, were notably reduced under this pharmacological intervention, emphasizing the efficacy of serotonergic modulation in enhancing resilience to proteotoxic stress (fig. S7F and G).

Next, we sought to investigate the sensitivity of the assay to detect the contribution of *mod-5* into whole-organism response to HS and its utility for target identification. When exposed to 60 min of HS, *mod-5* ablated animals demonstrated a notably enhanced resilience to HS compared to WT animals (Fig. 6). Particularly, *mod-5* ablation exhibited a protective effect immediately after HS (Fig. 6A and B). As WT, *mod-5* ablated animals restored their motor activity to baseline conditions within 15 hours post-HS (Fig. 6C), however, their recovery initiation occurred much earlier, beginning at 60 min post-HS, compared to 320-360 min in WT (Fig. 6A). These findings are corroborated by the total activity during recovery and T50 values (Fig. 6D, E, respectively). In contrast, activity of *mod-5* animals did not change upon CIT treatment at the most effective concentration identified in this assay (Fig. 6F-J), suggesting drug and assay specificity. Taking this together, these results confirm *mod-5* as a proteostasis suppressor gene, highlighting the potential of this novel method to validate candidate proteostasis-related drug targets or identify new ones through genetic or pharmacological approaches.

**Figure 6.**
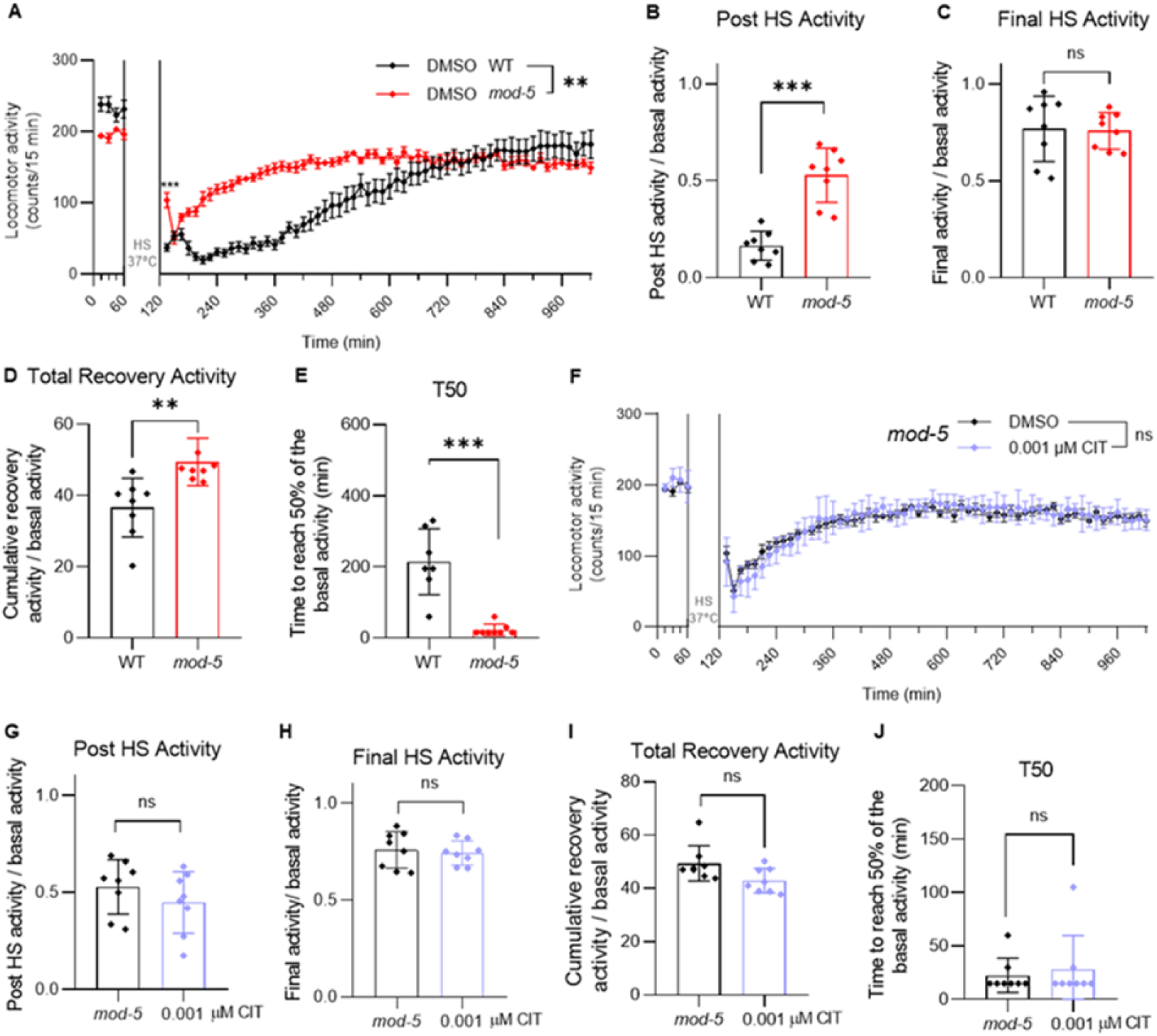
Differential responses to HS in *mod-5* mutant and WT *C. elegans.* **(A)** Illustrate the immediate response of *mod-5* mutants and wild-type (WT) strains when exposed to 60 minutes of HS at 37 °C. Statistical significance determined by Two-way RM ANOVA, Tukey’s multiple comparisons test. **(B)** Post-HS activity- assessed for 15 min after the removal of the HS stimulus-divided by the mean of the basal activity of each experimental group. **(C)** Total recovery activity of the animals measured over a 15-hour recovery period post-HS (normalized to basal activity). **(D)** Time to achieve 50% recovery of the basal activity after HS (T50) of WT and *mod-5* mutant animals. **(E)** Final recovery activity of WT and *mod-5* nematodes was measured during the last 60 min of the assay and normalized to the corresponding basal activity. (**B-E**) This finding is supported by at least three independent trials. Each biological experiment included eight technical replicates (N=400) for each concentration. *P* values were calculated using t-test. The data are presented as mean ± SD. **(F)** Investigation of MOD-5’s role in citalopram-mediated protection against HS-induced stress. Statistical significance determined by Two-way RM ANOVA, Tukey’s multiple comparisons test. **(G-J)** Analysis of post-HS activity, cumulative recovery activity, T50, and final recovery levels. This finding is supported by at least three independent trials. Each biological experiment included eight technical replicates (N=400) for each condition. *P* values were calculated using t-test. The data are presented as mean ± SD.

### Assay protocol adaptation reveals increased thermotolerance of animals treated with CCI-779 and citalopram

Thermotolerance assays are often used to investigate whole-organism responses to heat stress and to establish signaling pathways underlying these responses (*43*, *64*, *65*). However, these assays are time-consuming and highly dependent on the experimenter. Other limitations include (1) the fact that survival is frequently used as a readout, which implies a discontinuous assessment of the effects of elevated temperatures, and (2) that temperature shifts often occur during the scoring periods. Here, we combined a programable temperature-controlled incubator with the automated worm activity tracking system and used animals’ activity counts as a measure of thermotolerance. In this variant of PRO-FitS, synchronized *C. elegans* eggs were incubated with food for three days. Baseline motor activity was recorded at 20 °C for 60 min (basal activity), after which temperature started to rise until the desired final temperature of 33 °C, which was maintained throughout the assay. Motor activity data was collected continuously (values every 5 min or more), providing a temporal profile of the impact of temperature increase on animals’ motor activity (Fig. 7A). Importantly from the point-of-view of assay development, this experimental setup is amenable to easy modification, allowing changes in temperature levels and exposure time lengths, permitting protocol customization according to the research question. With the PRO-FitS assay variant, we were also able to detect an effect of CCI-779 treatment, which significantly sustained motor activity under elevated temperature conditions, for 90 min at 33°C, when compared to their untreated counterparts (Fig. 7B). Similar results were obtained with CIT administration, with CIT-treated nematodes exhibiting significantly increased locomotor activity (during 120 min), compared to those receiving vehicle-control treatment (Fig. 7C). This enhancement in motor activity upon sustained heat stress further advocates the neuroprotective effects previously described for both compounds (*48*, *49*, *59*, *62*, *66*). We further validated this assay using a tauopathy model of *C. elegans*. Hyperphosphorylated, insoluble, and filamentous tau is a hallmark of many NDs, collectively known as tauopathies (*67*). For our experiments, we selected a transgenic *C. elegans* model of the human tauopathy FTDP-17 (*68–70*), which expresses pan-neuronal full-length human mutant Tau V337M. Mutant Tau (mTau) pan-neuronal expression causes locomotion defects, as extensively described previously in several motor behavior paradigms. Using PRO-FitS, we found that CCI-779 treatment significantly enhanced motor activity at 33 °C of mTau animals (Fig. 7D). Together, our results pave the way for the use of PRO-FitS (as well as possible assay variants) for testing novel proprietary small molecules, as well as to repurpose or reposition other compounds with previous known applications as therapeutic approaches for disorders of protein dyshomeostasis.

**Figure 7.**
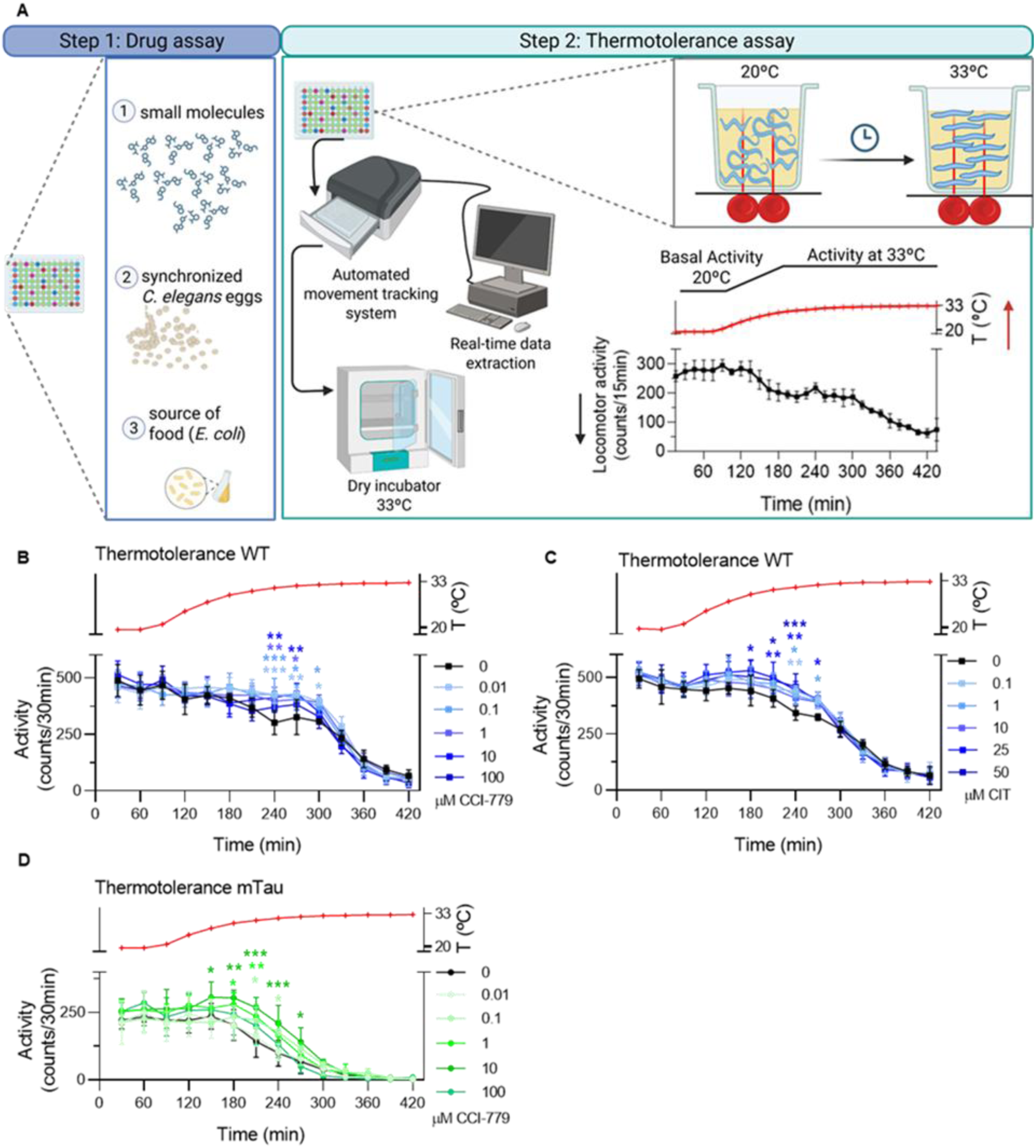
Assessment of motor activity resilience to elevated temperature of WT and mutant Tau-expressing *C. elegans* upon drug treatment, as a measure of thermotolerance. **(A)** Schematic representation of the experimental protocol. Synchronized *C. elegans* eggs were incubated with citalopram or CCI-779, and *E. coli* OP50 as a nutritional source in a 96-well plate (step 1). On the third day, nematodes were placed in a WMicrotracker system inside a dry incubator. Motor activity was first measured at 20 °C, followed by a gradual increase to 33 °C while movement was continuously monitored (step 2). **(B-C)** Effects of CCI-779 and citalopram on the thermotolerance of WT *C. elegans*. **(D)** Thermal resilience in the tauopathy model strain CK10 (*C. elegans* expressing human mutant tau), treated with CCI-779. Data were analyzed using two-way RM ANOVA with Tukey’s multiple comparisons test. Each condition was tested in at least three independent experiments, with 8 technical replicates (N=400 animals) per condition. Data are presented as mean ± SEM.

## Discussion

Disrupted proteostasis is a hallmark of aging and a central feature in the pathogenesis of distinct neurodegenerative diseases (NDs). Despite ongoing research efforts, the identification of proteostasis modulators and a deeper understanding of their temporal and functional dynamics remains incomplete. Given the urgent medical need for effective therapeutic approaches for NDs, we presented a novel phenotypic assay to streamline the identification of compounds with proteostasis-enhancing capacity, with the potential for mitigating the proteotoxic stress associated with neurodegenerative disorders. Using *C. elegans* as the *in vivo* platform, we established a large-scale screening method that provides automated and quantitative real-time readouts of motor activity as an indicator of proteostasis disruption induced by acute heat stress (Fig. 1). This dynamic assay captures the effects of proteostasis modulation within the context of an entire organism under induced stress conditions.

Several high-throughput methods to identify proteostasis enhancers have been previously described, some of which focused on specific branches of the proteostasis network. For instance, cell-based screening using fluorescent or luminescent reporters under the regulation of HS inducible promoters identified molecules capable of activating HSF1 (*71*, *72*). Similar cell-based reporter approaches have been used to identify chemical and genetic inductors of autophagy (*73–75*). Proteasome activity probes (*76*) or reporters (*77*) were employed to find Ubiquitin-Proteasome System activators. While these approaches provided valuable insights by screening chemical and genetic modulators of proteostasis, they fail to fully account for systemic, organism-wide effects as they do not include a functional nervous system or allow behavioral assessments. Our approach, by contrast, offered a more dynamic and integrated perspective of proteostasis regulation, reflecting the network of biological processes involved at multiple levels of organization. Additional whole-organism screenings have also been described. For example, genome-wide RNAi screenings were used with different folding sensors, aggregation-prone proteins, or reporter strains in *C. elegans* (*78–80*). Drug screenings were also reported using the same paradigms (*81–83*). Moreover, several screenings have been based on the assessment of specific behaviors affected in ND models; among others (*84*, *85*), we previously described a small-molecule screening that used the motor phenotype of a *C. elegans* model of MJD/SCA3 as a read-out (*58*, *62*).

Despite all the advances obtained with these screening methodologies, an assay allowing unbiased identification of proteostasis enhancers with a broader applicability was still missing. With this phenotypic assay, we demonstrated that acute HS effectively induces protein misfolding, thereby challenging proteostasis and temporarily impairing motor performance in *C. elegans*. In addition, the optimized HS protocol provided a reliable means to study the effects of proteotoxic stress, yielding consistent and reproducible results. Importantly, this approach enabled the longitudinal assessment of motor function with minimal influence of the experimenter, offering a more dynamic and objective evaluation compared to other thermorecovery and thermotolerance assays, which typically rely on measures such as survival rates, which are limited, and protein aggregation changes, which can be subjective (*32*). While a previous study using a heat stress paradigm in *C. elegans* to identify genetic enhancers of proteostasis has been described (*86*), the PRO-FitS takes it a step further and assesses how the proteotoxic stress impacts both behavior and protein aggregation. This methodology provides a clear and comprehensive view on how proteotoxic stress impacts overall organismal function. Yet, the proteotoxic insult induced via HS in PROFitS is inherently acute and thus very different from the prolonged, chronic stress characteristic of NDs, which represents a limitation for this assay (*87–89*). This acute thermal stress may lead to aggregation of proteins that are particularly sensitive to heat, rather than replicating the selective, cell-type-specific protein misfolding observed in NDs. HS was used as a proxy for proteostasis failure, effectively denaturing and misfolding a broad range of proteins simultaneously. This approach induces an overwhelming stress to the proteostasis machinery, partially mimicking the later stages of NDs. Additionally, by utilizing a general proteotoxic stress triggered by HS rather than the expression of a specific protein or genetic environment, it becomes possible to identify compounds that enhance proteostasis without being context-dependent on specific proteins associated with NDs. This is particularly relevant given that many ND share common pathological events related to the disruption of proteostasis machinery (*5*). As a proof-of-concept for the assay’s validity, proteostasis enhancing treatments that have already shown efficacy in ND context in higher organisms, (e.g. temsirolimus and citalopram), as well as genetic modifications with similar effects (MOD-5/SERT KO), promoted a faster recovery of animals to the thermal insult (*49*, *62*, *90–92*). The assay was also applied to *C. elegans* models of NDs, such as MJD/SCA3 and tauopathy, providing similar results. These results strongly corroborate the high potential of the PRO-FitS platform for the identification of effective proteostasis modulators, valuable for preclinical and clinical development.

Besides pharmacological screenings, PROFitS offer a versatile platform that can be adapted for a wide range of applications. After identification of a pharmacophore, structure-activity relationship studies can be streamlined, systematically testing different chemical derivatives with modifications in functional groups to assess their effects on biological activity, optimizing potency, selectivity and bioavailability of new therapeutic leads. Target dependence and mechanism of action can also be elucidated by employing genetic mutants or RNAi-mediated gene silencing. Since this assay utilizes miniaturized liquid cultures in parallel, *C. elegans* bacterial diet can be modified to explore the impact of different microbiota-derived metabolites on proteostasis. The gut microbiome is increasingly recognized as a key regulator for host protein homeostasis and ND disease progression (*93–95*). Identifying bacteria that produce metabolites with proteostasis-enhancing properties or uncovering microbial-derived factors that induce proteotoxic stress or impair cellular resilience is crucial for understanding the complex interplay between the gut microbiome and host health (*94*, *96–99*). Identification of agents that exacerbate proteotoxic stress, which can be useful as anticancer therapy or to identify environmental risks, can also be a powerful outcome of this assay (*100*, *101*). The flexibility of this phenotypic assay makes it a powerful tool to study proteostasis from multiple perspectives and broadens its application in ND research and drug discovery.

Another key advantage of this approach is compatibility with other high-throughput systems using the same 96-well plate format. For example, automated imaging platforms, capable of capturing fluorescence across the entire plate can be seamlessly integrated with the WMicroTracker system (*102*). By employing fluorescent reporter strains, researchers can sequentially assess the effects of treatments on gene expression while measuring motor behavior within the same experimental setup. This dual-readout approach provides complementary insights into both molecular and functional outcomes, enabling a more comprehensive evaluation of treatment effects. This accelerates the screening process and allows for a more efficient use of resources by reducing the number of required experiments, minimizing reagent consumption, and maximizing data output from a single assay. In the same line of resource optimization, the implementation of this phenotypic assay in *C. elegans* significantly contributes to the 3Rs principle—Replacement, Reduction, and Refinement—by providing an efficient and scalable alternative to early-stage rodent testing. Allowing researchers to screen thousands of small molecules, perform structure-activity relationship studies and target dependency minimizes the need for preliminary studies in mammalian animal models. While the phylogenetic distance between *C. elegans* and humans may raise concerns regarding the immediate translational potential, it is well-established that key proteostasis network modules, such as chaperones, autophagy and proteasome components, and organelle stress responses, are highly conserved between nematodes and mammalians (*103–105*). Therefore, *C. elegans* is a valuable model for first-line identification of small molecule proteostasis enhancers, possibly translatable to human disease contexts.

In conclusion, PROFitS provide a robust and thoroughly validated platform for investigating proteostasis modulation. We demonstrate that using motor behavior as a proxy for the state of the proteome is an effective approach. Taking advantage of this, and by inducing a proteotoxic insult in *C. elegans* and following motor behavior recovery, allows the assessment of small molecules that may enhance cellular resilience to such stress. Furthermore, we establish its relevance in the context of NDs. The assay’s scalability, adaptability, and compatibility with complementary high-throughput techniques further enhance its utility in drug discovery and development, as well as mechanistic studies. Ultimately, the implementation of this method is highly valuable for the identification of novel therapeutic strategies for NDs, while also expanding our understanding of proteostasis regulation at the whole-organism level.

## Materials and Methods

### *Caenorhabditis elegans* strains and maintenance

*C. elegans* were maintained using standard methods at 20 °C on nematode growth medium (NGM) plates, seeded with the *Escherichia coli* (*E. coli*) OP50 strain. Bacteria were grown overnight at 37 °C at 180 rpm in Luria Broth (LB) medium (*106*). *C. elegans* strains used in this work are the following: wild-type (WT, N2 Bristol), MT8944 *mod-5*(n882) I (*63*), DCD23 *uqIs5*[lbp-2p::lbp-2::tagRFP] (*107*), ZG120 *unc-119*(*ed3*) III; *iaIs7* IV (*108*) and ZG31 *hif-1*(*ia04*) V (*45*), all obtained from the *Caenorhabditis Genetics Center* (CGC), University of Minnesota. Additional strains included MJD-related strains (AT3Q75 (*rmls237*[PF25B3.3::*AT3v1-1q75::*YFP]) and AT3Q130 (*rmls263*(PF25B3.3::*AT3v1-1q130::*YFP) II)), previously generated by us (*58*) and the *C. elegans* model of FTDP-17 (*bkIs10*[Paex-3::h4R1NTauV337M;*Pmyo-2*::GFP]) (*68*), kindly provided by Dr. Brian Kraemer.

### Compounds

Compounds were obtained from the commercial vendors indicated below and were used without further purification: temsirolimus (CCI-779) CAS 162635-04-3 (LC laboratories) and citalopram hydrobromide CAS 59729-32-7 (Kemprotec Ltd.).

### Toxicity assessment of CCI-779

The *C. elegans* Bristol strain N2 was used to evaluate the toxicity of CCI-779, whereas citalopram toxicity was assessed elsewhere (*62*). The assay was performed in 96-well plates, in liquid culture, as already described (*62*). Briefly, each well contained a final volume of 60 μL, comprising 15 μL of egg suspension (20–25 eggs), 25 μL of CCI-779 at 9 different concentrations (0.000001 to 100 µM) and 20 μL of OP50 bacteria at an OD 595 nm of 0.6–0.7. The OP50 bacteria were previously grown overnight at 37 °C and 180 rpm in LB, pelleted by centrifugation, inactivated by 3 cycles of freeze/thawing, frozen at −80 °C and then resuspended in complete S-medium. Worms were grown with continuous shaking at 180 rpm at 20 °C (Shell Lab incubator shaker) for 7 days. The effect of compounds on *C. elegans* physiology was monitored by evaluating the rate at which the *E. coli* food suspension was consumed, as a readout for *C. elegans* growth, survival, or fecundity. The absorbance (OD_595nm_) was measured daily (NanoQuant Plate™-Tecan). DMSO 1% (vehicle, non-toxic) and DMSO 5% (toxic condition) were used as controls.

### Drug assays in liquid medium

Drug assays were performed in a 96-well plate format, in liquid culture, as previously described (*42*, *62*), with some modifications, that included adjusting the volumes to be compatible with the WMicrotracker system. Age-synchronized populations of *C. elegans* were prepared by treating gravid adults with an alkaline hypochlorite solution (0.5M NaOH, ∼2.6% NaClO) for approximately 5 minutes. Following treatment, the animals were washed twice with M9 buffer (22 mM KH**_2_**PO**_4_**, 22 mM NaH**_2_**PO**_4_**, 85 mM NaCl, 1 mM MgSO**_4_**) and subsequently resuspended in S-medium to achieve a final concentration of 50 eggs per well. OP50 *Escherichia coli* bacteria were cultured overnight at 37 °C and 180 rpm in LB medium. The bacteria were then pelleted by centrifugation, inactivated through three cycles of freeze-thawing, and stored at −80°C. Prior to use, the inactivated bacteria were resuspended in S-medium supplemented with cholesterol, streptomycin/penicillin, and nystatin (Sigma) to an optical density (OD) at 595 nm of 0.5-0.6, measured using a NanoQuant Plate™ (Tecan). Compounds were initially prepared as 100 mM stock solution in 100% dimethyl sulfoxide (DMSO, Sigma) and stored at −20 °C. On the day of the assay, each compound was further diluted in Milli-Q purified H**_2_**O to achieve final concentrations ranging from 100 µM to 1 pM (CCI-779 and citalopram, CIT) in a solution containing 1% DMSO (the drug-vehicle concentration was kept constant across the different experimental conditions). This dilution strategy was employed to minimize any solvent-specific toxic effects that could potentially impact the nematodes negatively. Each well of a 96-well plate contained 55 µL of inactivated bacteria suspension, 35 µL of animal suspension, and 10 µL of drug or drug-vehicle. Plates were kept under continuous shaking at 180 revolutions per minute (rpm) at 20 °C (Shel Lab) for three days. This environmental setting was chosen to ensure optimal growth and development conditions for the nematodes throughout drug exposure time.

### *C. elegans* heat shock protocol

On the third day post animal synchronization, the 96-well plate containing *C. elegans* was positioned on the WMicrotracker (*41*) system. This setup was utilized to measure the basal locomotor activity of the nematodes. The activity was recorded for 60 minutes, with data counts being registered every 15 minutes to establish a baseline prior to the heat shock (HS) treatment. Following the basal activity measurement, the plate was transferred to a temperature-controlled dry incubator set at 37°C. The plate was subjected to agitation at a speed of 180 rpm to ensure temperature distribution and to minimize temperature gradients within the incubator. The HS was induced during 30, 45, 60, 90, or 120 minutes, during the optimization steps and then established to be 60 minutes. Immediately following the HS, the plates were quickly relocated back to the WMicrotracker device to continue monitoring the post-HS recovery activity of the animals. The assessment of the post-HS recovery activity took 15 hours, or the indicated durations. Continuous monitoring allowed for detailed analysis of the kinetics of motor activity recovery.

### Thermotolerance Assay

On the third day following animal synchronization, a 96-well plate containing *C. elegans* was placed in the WMicrotracker system to monitor motor activity. The device was then placed inside a temperature-controlled incubator initially set to 20 °C. The animals’ activity was recorded at this temperature for 1 hour to establish a baseline under stable thermal conditions. After this period, the temperature was gradually increased up to 33 °C. Locomotor activity was continuously recorded throughout the temperature ramp-up and for an additional 6 hours after the onset of temperature increase.

### pH measurement with and without shaking

The pH of the culture medium was assessed under two conditions: with and without shaking. The liquid-based assay was performed as described above. On day 3 of the experiment, one plate was maintained under continuous shaking (180 rpm; 20 °C), and another plate was placed in the WMicrotracker device without shaking for 12 hours at 20 °C. After this period, the pH of the medium in both plates was measured using Labbox Universal Indicator Strips (Labbox). The strips were photographed, and the colors were compared to the reference color scale provided by the manufacturer to determine the corresponding pH values.

### Transcriptional reporter assay for hypoxia induction

Age-synchronized *C. elegans* ZG120 reporter strain were cultured in liquid medium under standard conditions. On day 3 after hatching, one plate was maintained under continuous shaking (180 rpm; 20 °C), while the other plate was placed without shaking for 24 hours to evaluate whether lack of agitation could induce hypoxia-related gene expression. On day 4, worms were anesthetized with 2 mM sodium azide (Sigma, St. Louis, MO) and mounted on 3% agarose pads. Animals were oriented using an eyelash. Excess anesthetic was removed, and worms were covered with a cover slip and sealed with 3% agarose. GFP fluorescence images were acquired using an Olympus IX81 inverted fluorescence microscope with an Olympus LUCPlanFL N 10x objective, NA 0.30 Ph1, coupled with an Olympus XM 10 (CCD gray scale, 14bit) camera. Imaging parameters were set based on the control condition (with agitation) and maintained constant across all samples. Fluorescence intensity was measured using Fiji (ImageJ; National Institutes of Health, Bethesda, MD), normalized to the total area of each animal, and expressed relative to the mean fluorescence of the control group (shaking condition). The experiment was independently performed four times, analyzing at least ten animals per condition in each replicate.

### Assessment of hypoxia induction

Age-synchronized eggs of ZG31 [*hif-1* (*ia04*) V] strain were distributed into 96-well plates at a density of approximately 50 eggs per well. Worms were grown in liquid culture medium under conditions described above. On day 3 post-synchronization, the plate was placed in the WMicrotracker device, and locomotor activity was recorded automatically over a 15-hour period under static (non-agitated) conditions. Eight wells were analyzed, totaling approximately 400 animals per assay. Two biological replicates were performed.

### Filter retardation assay

A drug assay was conducted as described above. After three days, the animals (from the entire 96-well plate) were collected in pre-, post-, 2h- and 4h-post-HS, washed 3x with M9 and flash frozen in liquid nitrogen. For protein extraction, the worms were disrupted in a non-denaturating buffer using glass beads (Sigma), in a FastPrep®-24 (MP Biomedicals) device, as described in detail in (7). Protein concentration was quantified using the BCA assay and 100 μg of total protein were loaded into a cellulose acetate membrane mounted in a slot blot apparatus (Bio-Rad). Amyloid fibrils and fibrillar oligomers retained on the membrane were detected with rabbit anti-OC (1:100, Millipore, Cat# AB2283) and chemiluminescence (ECL Radiance, Azure Biosystems). Chemiluminescence signals were captured with the Sapphire Biomolecular Imager (Azure Biosystems), and quantified using AzureSpot Analysis Software (Azure Biosystems), according to the manufacturer’s guidelines. Separate gels were loaded with the same samples (10 µg) and stained with AzureRed (Azure Biosystems), according to the manufacturer’s instructions, to obtain total protein loading for each sample. Amyloid fibrils and fibrillar oligomers signal in the membrane were normalized to the signal of total protein AzureRed-stained gel.

### Confocal imaging for quantification of aggregation *in vivo*

A synchronized population of approximately 500 *C. elegans* expressing LBP-2::TagRFP (strain DCD23) was obtained by egg laying on NGM agar plates seeded with *E. coli* OP50. Animals were maintained and monitored from day 4 to day 14 after hatching to assess protein aggregation levels by confocal microscopy at two-day intervals. After day 4, animals were transferred every two days to fresh NGM-seeded plates to prevent interference from progeny. LBP-2::TagRFP animals grown in NGM-seeded plates until day 6 after hatching were submitted to HS (60 min; 37 °C). The profile of LBP-2 aggregation was assessed by confocal microscopy at baseline (pre-HS) and at 2-, 4-, and 12-hours post-HS. For drug treatment, LPB2::TagRFP animals were exposed from the egg stage until day 4 after hatching to either vehicle control (1% DMSO) or CCI-779 at a concentration of 100 µM dissolved in inactivated bacteria. On day 4, animals from each treatment group were subjected to HS at 37 °C for 60 min. Imaging was performed at the following timepoints: pre-HS, immediately after HS (post-HS) and 2 h post-HS. For confocal dynamic imaging, live animals were immobilized using 10 mM levamisole (Sigma). Worms were mounted on a 3% agarose pad. All images were captured using an Olympus Confocal Laser Scanning Microscope FV3000, utilizing an Olympus UPlaPon 60x OHR objective, NA 1.50. Imaging was conducted using a 561 nm laser line to excite red fluorescent protein (RFP). The pinhole was set to 1.0 Airy unit to optimize optical slice thickness. Z-series scans were acquired at intervals of approximately 0.75 μm along the Z-axis. For quantification of aggregates, Z-projections of the scanned images were generated using FV10-ASW 4.2 viewer software (Olympus). Aggregates were specifically quantified within defined anatomical regions of the worms, namely the head region, anterior bulb, and tail region, as detailed in the figure legend associated with the results. Based on the quantification results, animals were categorized into three distinct groups: animals with no visible puncta, animals with up to ten puncta, and animals with more than ten puncta.

### Motor behavior assay in the *C. elegans* model of MJD/SCA3

The motor phenotype of the MJD/SCA3 model was evaluated using a motility assay that evaluates crawling, as previously described (*62*). At day 4 after hatching, the animals were removed from the liquid medium to unseeded NGM plates, for 45 min. Five to ten animals were picked onto the center of a plate, homogenously seeded in all its surface and equilibrated at 20 °C. After 1 min, animals remaining inside a drawn 1 cm circle were scored as locomotion defective. A total of ∼50 animals were scored per assay, and at least three independent assays were performed for each condition.

### Trashing assay

The thrashing assay was conducted following the same experimental setup as previously described for the drug assay and HS induction. On the assay day, 10 μL drops of M9 buffer were carefully placed on clean glass slides. Special care was taken to ensure that the buffer drops remained well-separated and did not spread, thereby maintaining their integrity as discrete droplets suitable for the assay. Individual worms were picked using a standard platinum wire worm picker and gently transferred to each M9 buffer drop on the prepared slides. Worms were allowed to acclimate and swim freely for 30 seconds. This brief acclimatization period was critical to allow the worms to recover from the mechanical stress of transfer and adapt to the new medium, ensuring consistent behavior during filming. Each worm was filmed for a period of one minute using an Olympus PD72 digital camera attached to an Olympus SZX16 stereomicroscope. The camera setup was adjusted to ensure that the entire M9 buffer drop remained within the field of view to continuously capture the worm’s activity. Adequate lighting and focus were carefully maintained throughout the recording process to capture high-quality video data of the worm’s thrashing movements. The number of thrashes performed by each worm was counted for one minute. Trash was defined as a worm movement that swings its head and/or tail to the same side. This metric was used to assess the neuromuscular activity and the impact of drug treatment on the locomotive behavior of the worms before and 0-, 2-, 4-, 6 and 24-hour post-HS. We monitored at least 10 worms for each condition in each independent assay.

### Statistical Analysis

A confidence interval of 95% was assumed for all statistical tests with significance levels of p < 0.05 (*), p < 0.01 (**) and p < 0.001 (***), respectively. All continuous variables are presented as the mean and error bars represent the standard error of the mean, unless stated otherwise. Statistical analyses were performed using GraphPad Prism 8.0.1 software. The continuous variables distributions were tested for normality using Shapiro-Wilk or Kolmogorov-Smirnov normality tests and having into account sample size and histogram plot distribution. Levene’s test was also performed to verify homogeneity of variances between groups. Outliers were removed based on the ROUT method with a Q value set at 1%. For analysis of aggregation, two-sided Fisher’s exact test (GraphPad Prism, v.8) was performed to analyze two aggregation categories (animals with no puncta versus animals >10 puncta) and chi-square test (GraphPad) for three categories (animals with no puncta versus animals 1-10 puncta versus animals >10 puncta). For the comparison of more than two means, a one-way analysis of variance (ANOVA) was used, with a follow-up Tukey’s post hoc test. For the comparison of means with one independent and one repeated measures factor, a mixed design ANOVA with Tukey’s multiple comparison adjustment was used. For the comparison of two means, the student’s t test was used. Additionally, effect sizes for the dosages of treatments were calculated to quantify the magnitude of the observed effects. Statistical reports for all experiments are provided in Supplementary Table S1.

## Acknowledgments

We are grateful to members of the Translational Neurogenetics team for sharing reagents, for critical analysis of the data and discussions on the manuscript. Thanks to Dr. Brian Kraemer, who kindly provided the *C. elegans* model of FTDP-17. We also thank Drs Antonela Baron and Sergio Simonetta (Phylumtech) for their support and technical advice regarding the use of the WMicrotracker device. We are also grateful to the Caenorhabditis Genetics Center (CGC), which is funded by the National Institutes of Health – National Center for Research Resources, for providing nematode strains.

## Supporting Information for

### Figures

**Fig. S1.**
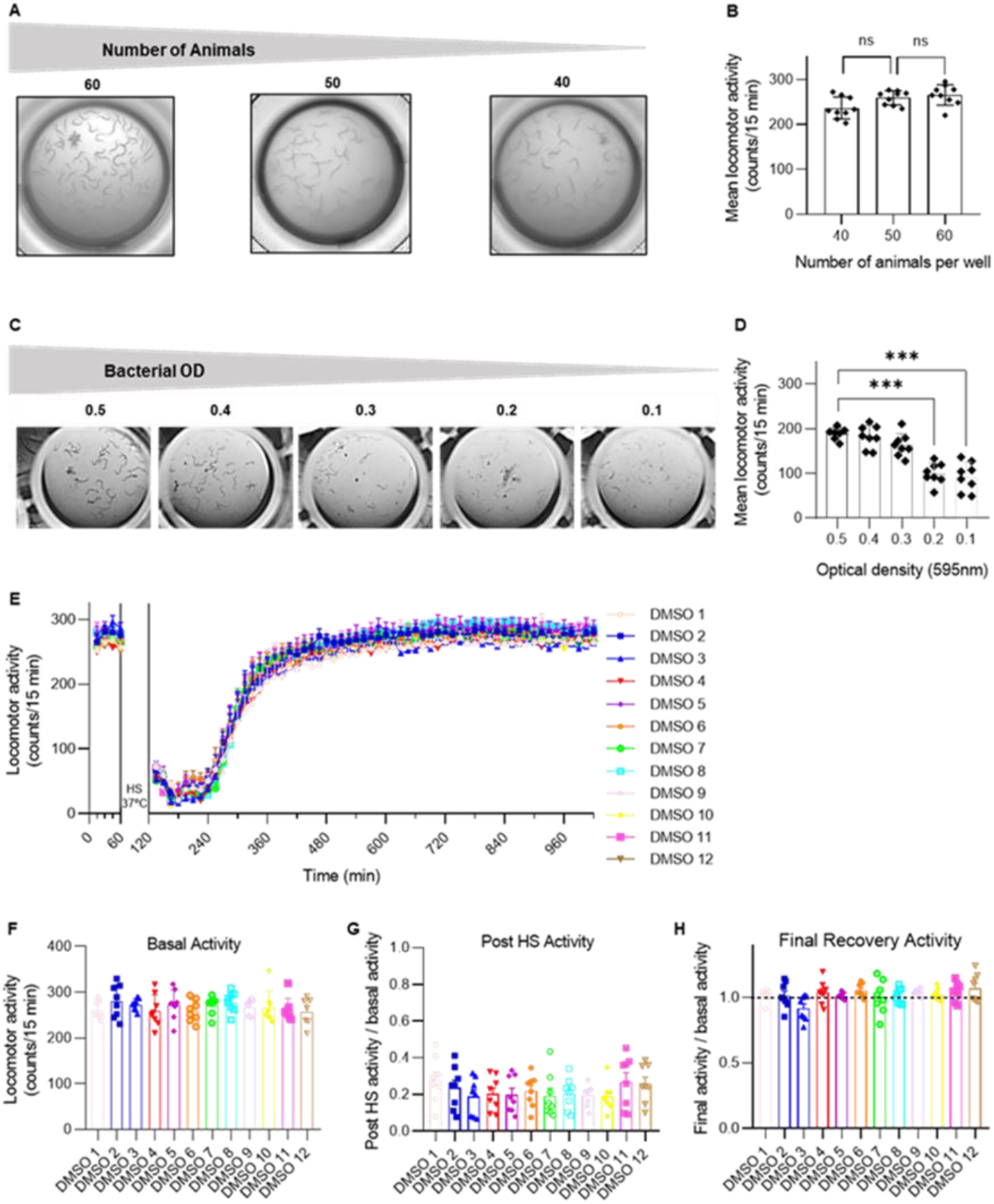
Impact of animal number, bacterial optical density and plate position on animals’ motor activity counts. **(A)** Representative images of distinct numbers of WT adult animals per well, ranging from 40 to 60 animals. **(B)** Locomotor activity in wells with 40, 50, and 60 animals, measured in the WMicrotracker. **(C)** Representative images of the impact of the different initial optical density (OD_595nm_) of bacteria on nematodes growth and development evaluated at day 3. **(D)** Effect of initial bacterial optical density on animals’ activity, quantified in the WMicrotracker at day 3. Data on the graphs represent mean ± standard deviation (SD) of motor activity in 8 wells per condition. *P* values (*P* < 0.001 (***)) were calculated using One-way ANOVA, Tukey’s multiple comparisons test. **(E)** Locomotor activity of animals growing under the same conditions (50 animals per well, initial bacterial OD of 0.5) analyzed across plate rows to assess any potential variability. Data on the graphs represent mean ± standard error of the mean (SEM) of motor activity in 8 wells (∼400 animals) per condition. *P* values (*P* >0.05 (ns)) were calculated using Mixed design ANOVA. (**F-H**) Basal, post-HS and final recovery activity counts analyzed across plate rows to assess variability. Data on the graphs represent mean ± SD of motor activity in 8 wells (∼400 animals) per condition. *P* values (*P* >0.05 (ns)) were calculated using One-way ANOVA, Tukey’s multiple comparisons test.

**Fig. S2.**
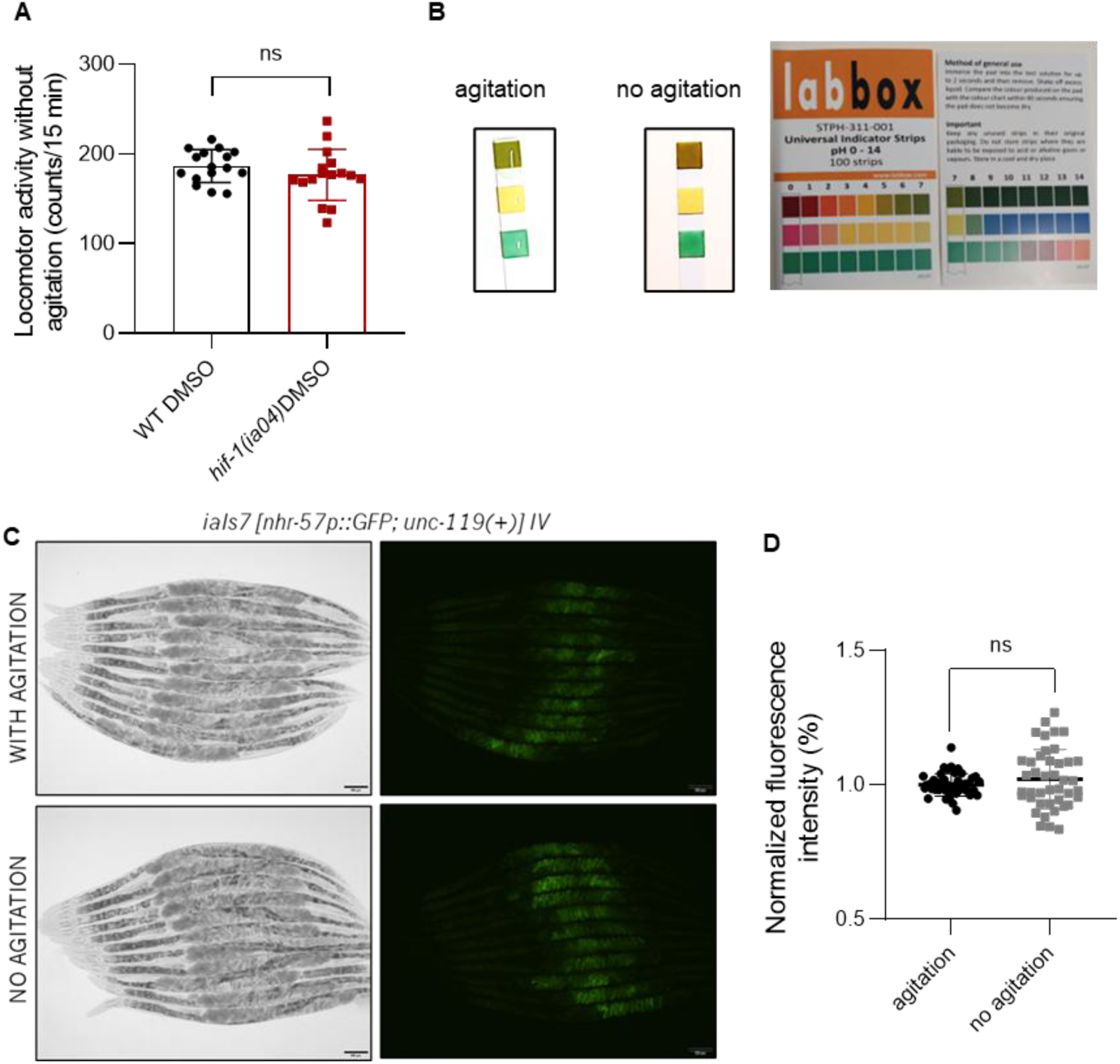
Impact of activity assessment conditions on the sensitivity of animals to hypoxia. **(A)** Assessment of the locomotor activity of the *C. elegans* strain ZG31 [*hif-1(ia04) V*] in the automated worm movement tracking system. Graph displays mean activity values ± SD of 8 wells (50 animals per well). The data represents mean values ± SD of 2 independent experiments with at least 400 worms analyzed in each assay. *P* > 0.05 (NS), t-test for comparison between the means of two groups. **(B)** pH levels of the growth media in plates maintained with agitation or kept in the plate reader without agitation. pH was measured with a universal pH indicator strip (LabBox). **(C)** Representative brightfield and fluorescence microscopic images of the reporter strain *nhr-57*p::GFP grown under the assay conditions with agitation or maintained in the plate reader without agitation (scale bar = 100 μm). **(D)** Relative fluorescence intensity of GFP proteins of the reporter strain *nhr-57*p::GFP grown under the assay conditions with or without agitation. The data represents mean values ± SD of 4 experiments with at least 10 worms analyzed in each assay. Data shown is normalized to the control condition (with agitation). *P* > 0.05 (NS), t-test for comparison between the means of two groups.

**Fig. S3.**
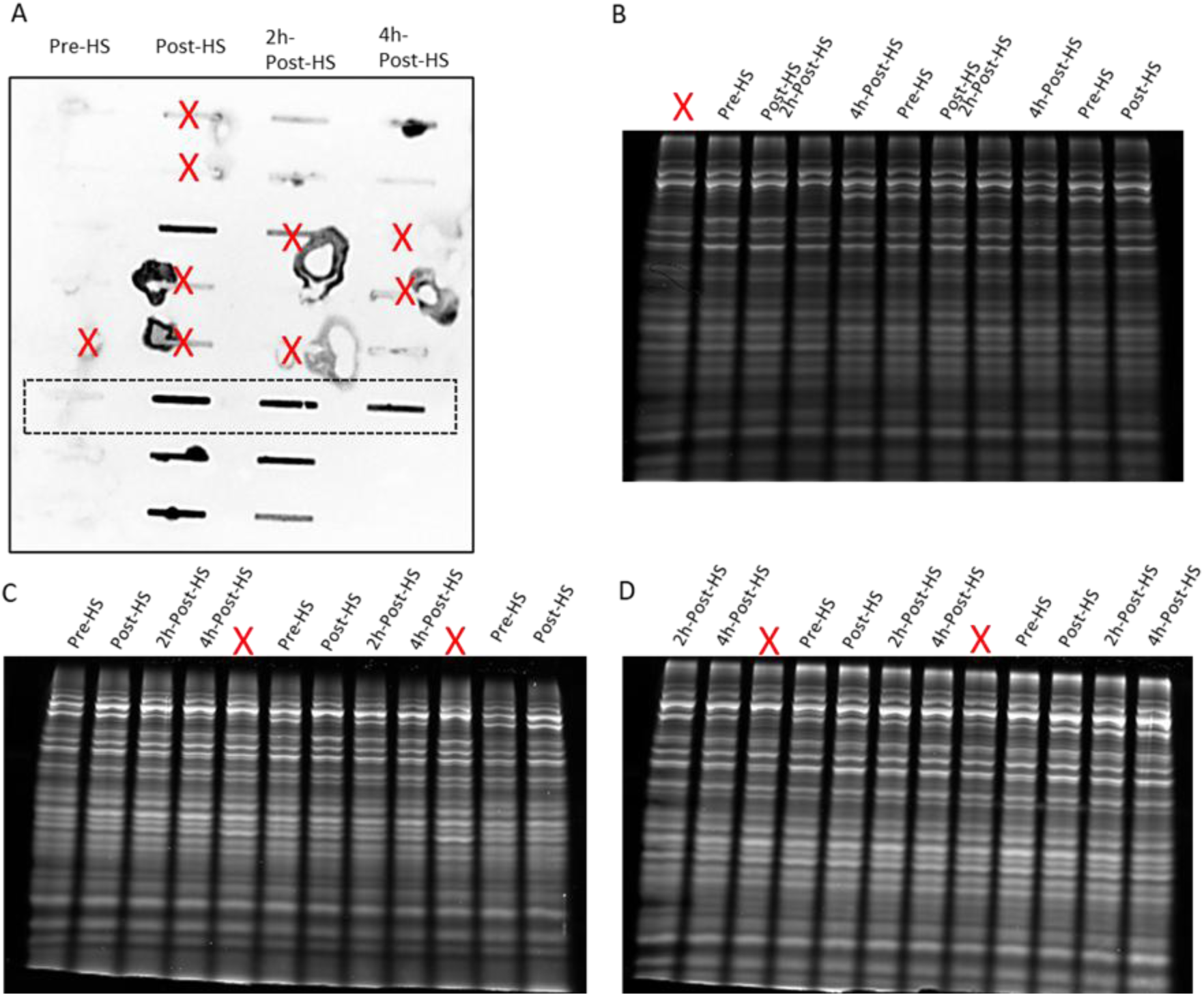
(Related with Fig. 3A) – **Original Western blot image presented in this study**. (**A**) Filter retardation assay blots of WT strain in Pre-, Post-, 2h- and 4h-Post-HS conditions. Some wells (indicated with an X) were eliminated from the analysis due to experimental reasons, namely the clogging of the filter, which prevented the samples from passing through the membrane efficiently. The highlighted rectangle (dashed, black) is shown in Fig. 3G. **(B-D)** Total protein gels of the samples of the filter retardation assay. Filter retardation assay blot densitometry values were normalized to total protein staining of these gels.

**Fig. S4.**
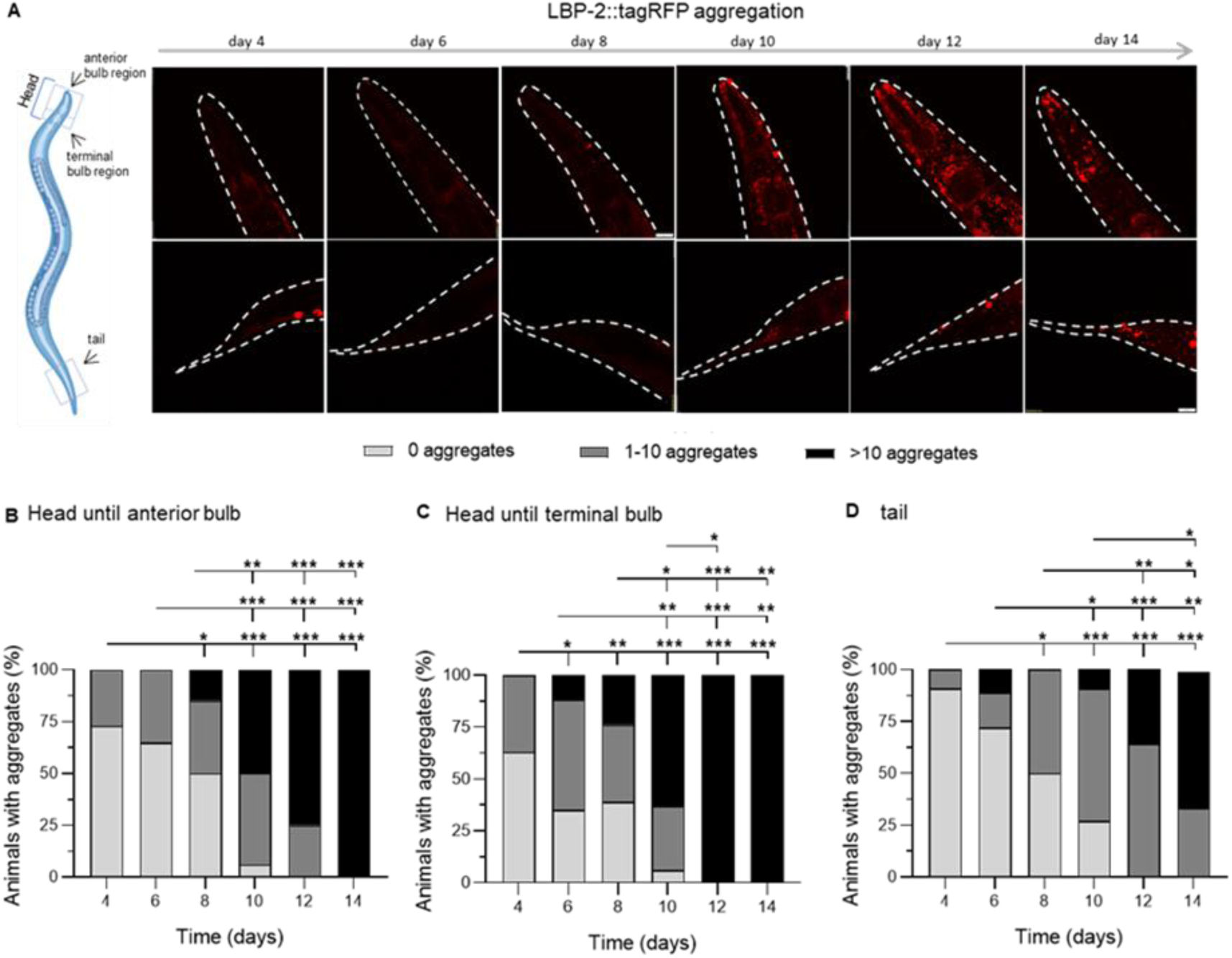
(Related with Fig. 3B) **Age-dependent extracellular aggregation of secreted LBP-2 in *C. elegans***. **(A)** Representative fluorescent microscopic images of age-dependent aggregation in LBP-2::tagRFP strain in specific regions of the body (illustrated in the *C. elegans* schematic representation) and timepoints for analysis. Scale bar = 20 µm. (**B-D**) Quantitative analyses of LBP-2::tagRFP aggregation in the head region until the anterior bulb **(B)**, head region until the terminal bulb **(C)**, and tail **(D)**. Each bar represents mean values ± SD of 3-4 independent experiments with at least 10 animals per condition in each. *P* values (*P* < 0.05) were calculated using the chi-square test and two-sided Fisher’s exact test.

**Fig. S5.**
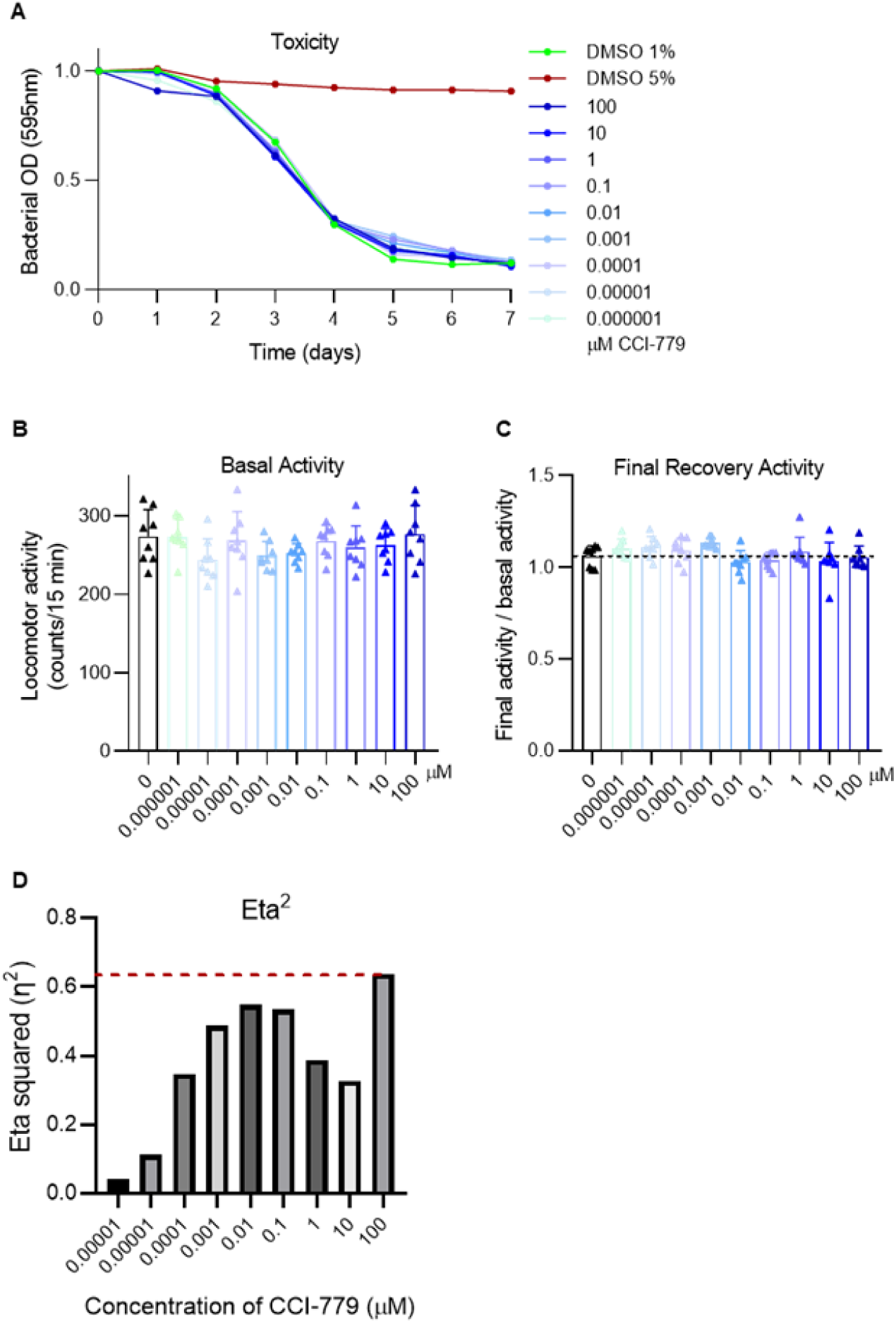
(related with Fig. 4) **Toxicity assessment, basal and final recovery activities of WT animals upon HS when treated with a range of concentrations of CCI-779. (A)** Daily bacterial consumption determined by optical density (OD) at 595 nm. Animals treated with CCI-779 were compared to control groups treated with 1% DMSO (non-toxic) and 5% DMSO (toxic), negative and positive controls of the assay, respectively. Measurements were taken in quintuplicate for each treatment condition (∼125 animals/group). *P* values were calculated using non-linear regression with four parameters: top, bottom, LogIC50 and hillslope. Data are presented as mean ± SD and values are normalized for day 0. **(B)** Basal locomotor activity of the animals was quantified in the automated plate reader during 60 min, following three days of treatment with CCI-779. **(C)** Final recovery activity of CCI-779-treated and untreated nematodes was measured during the last 60 min of the PRO-FiTS assay and normalized to the corresponding basal activity. **(D)** Effect size (η²) values from statistical analysis of the T50 parameter. The red dashed line indicates the η² value of the concentration with the highest effect size. **(C-D)** Data on the graphs represent mean ± SD values of the activity of 400 animals per condition. Data is representative of at least three independent trials. *P* values were calculated using One-way ANOVA, Tukey’s multiple comparisons test.

**Fig. S6.**
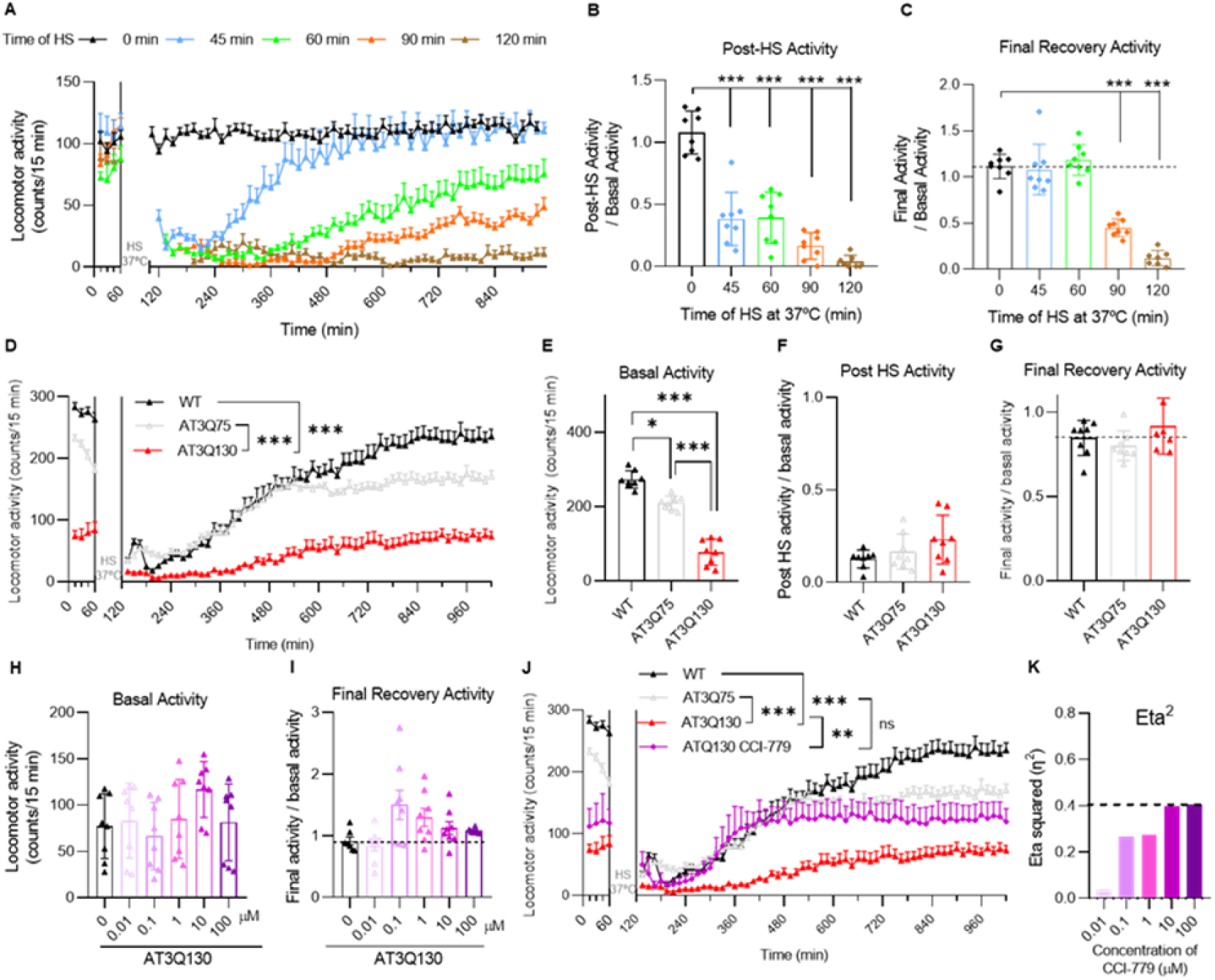
(related with Figure 5) Optimization of HS duration for assessing motor function recovery in a transgenic *C. elegans* model of Machado-Joseph disease (AT3Q130). (A) Measurement of locomotor activity following HS durations of 45, 60, 90, and 120 minutes. **(B)** Assessment of paralysis rates across different HS durations. **(C)** Final recovery activity was measured during the last 60 min of the assay and normalized to the corresponding basal activity. **(B-C)** Data on the graphs represent mean ± SD values of the activity of 400 animals per condition. Data is representative of at least three independent trials. *P* values were calculated using One-way ANOVA, Tukey’s multiple comparisons test. **(D)** Sensitivity of the assay in detecting genotype-specific differences in recovery following a 60-minute HS. Each biological experiment included eight technical replicates (N=400) for each condition. *P* values were calculated using Two-way RM ANOVA, Tukey’s multiple comparisons test. The data are presented as mean ± SD. **(E-G)** Analysis of baseline motor activity, immediate post-HS activity, and recovery to baseline levels 15-hour post-HS. Data on the graphs represent mean ± SD values of the activity of 400 animals per condition. Data is representative of at least three independent trials. *P* values were calculated using One-way ANOVA, Tukey’s multiple comparisons test. **(H)** Basal locomotor activity of the animals was quantified in the automated plate reader during 60 min, following three days of treatment with CCI-779. **(I)** Final recovery activity of CCI-779-treated and untreated nematodes was measured during the last 60 min of the assay and normalized to the corresponding basal activity. (H-I) *P* values were calculated using One-way ANOVA, Tukey’s multiple comparisons test. The data are presented as mean ± SD. **(J)** Recovery profile comparison of CCI-779-treated AT3Q130 animals (10 µM) and AT3Q75 animals during HS recovery. **(K)** Effect size (η²) values from statistical analysis of the T50 parameter. The red dashed line indicates the η² value of the concentration with the highest effect size. Each biological experiment included eight technical replicates (N=400) for each condition. *P* values were calculated using Two-way RM ANOVA, Tukey’s multiple comparisons test. The data are presented as mean ± standard deviation (SD).

**Fig. S7.**
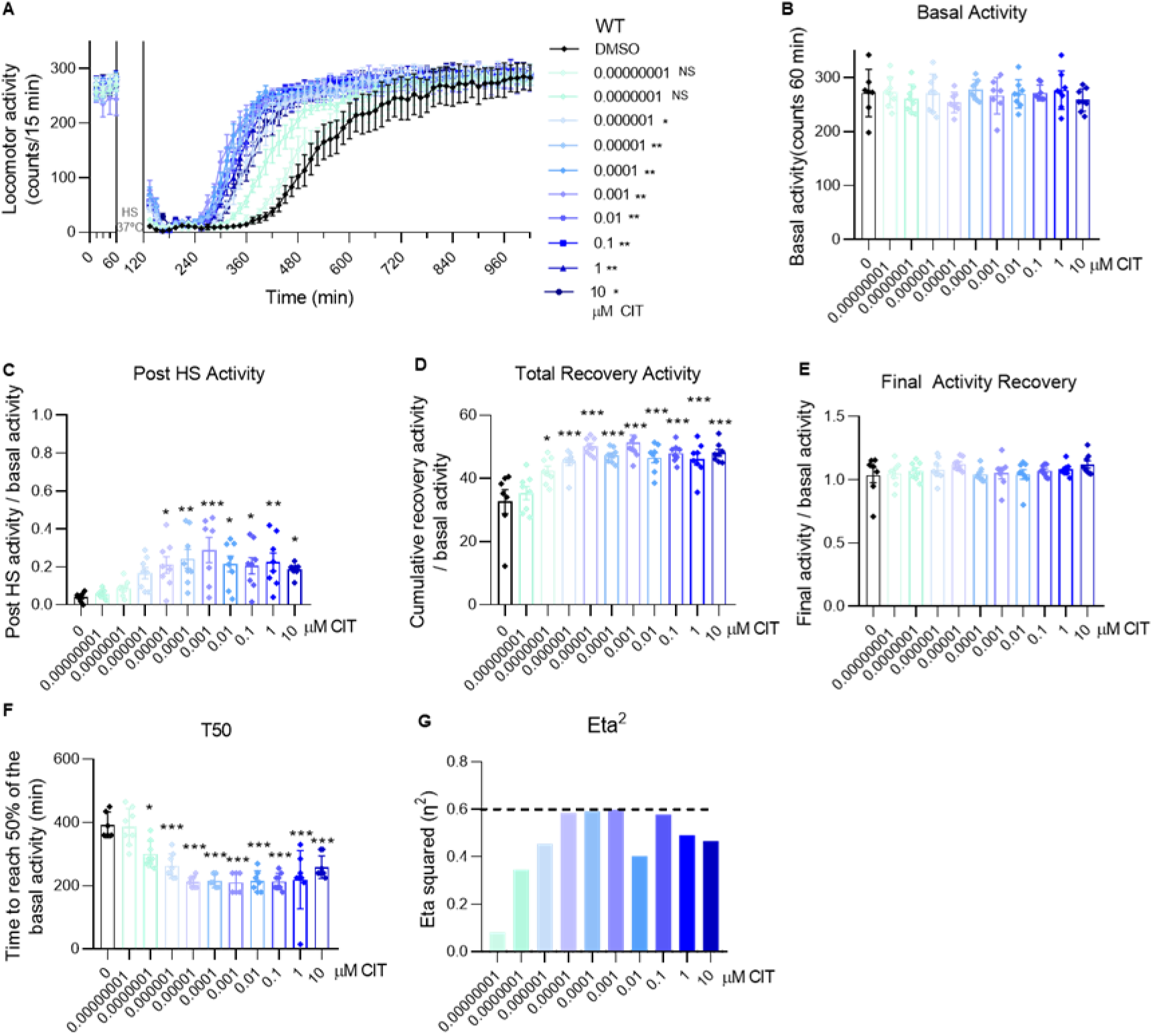
Assessment of citalopram (CIT) effects on HS-induced motor function in WT *C. elegans*. **(A)** Locomotor recovery following HS was monitored using the WMicrotracker system over a 15h period at 20°C. Data were analyzed using two-way RM ANOVA with Tukey’s multiple comparisons test. **(B)** Basal locomotor activity of the animals was quantified in the automated plate reader during 60 min, following three days of treatment with CIT. **(C)** Post-HS activity- assessed for 15 min after the removal of the HS stimulus- divided by the mean of the basal activity of each experimental group. **(D)** Total recovery activity of the animals measured over a 15-hour recovery period post-HS (normalized to basal activity). **(E)** Final recovery activity of CIT-treated and untreated nematodes was measured during the last 60 min of the assay and normalized to the corresponding basal activity. **(F)** Time to achieve 50% recovery of the basal activity after HS (T50) of CIT- and vehicle-treated animals. **(G)** Effect size (η²) values from statistical analysis of the T50 parameter. The black dashed line indicates the η² value of the concentration with the highest effect size. **(B-F)** Data on the graphs represent mean ± SD values of the activity of 400 animals per condition. Data is representative of at least three independent trials. *P* values were calculated using One-way ANOVA, Tukey’s multiple comparisons test.

### Tables

**Table S1.**
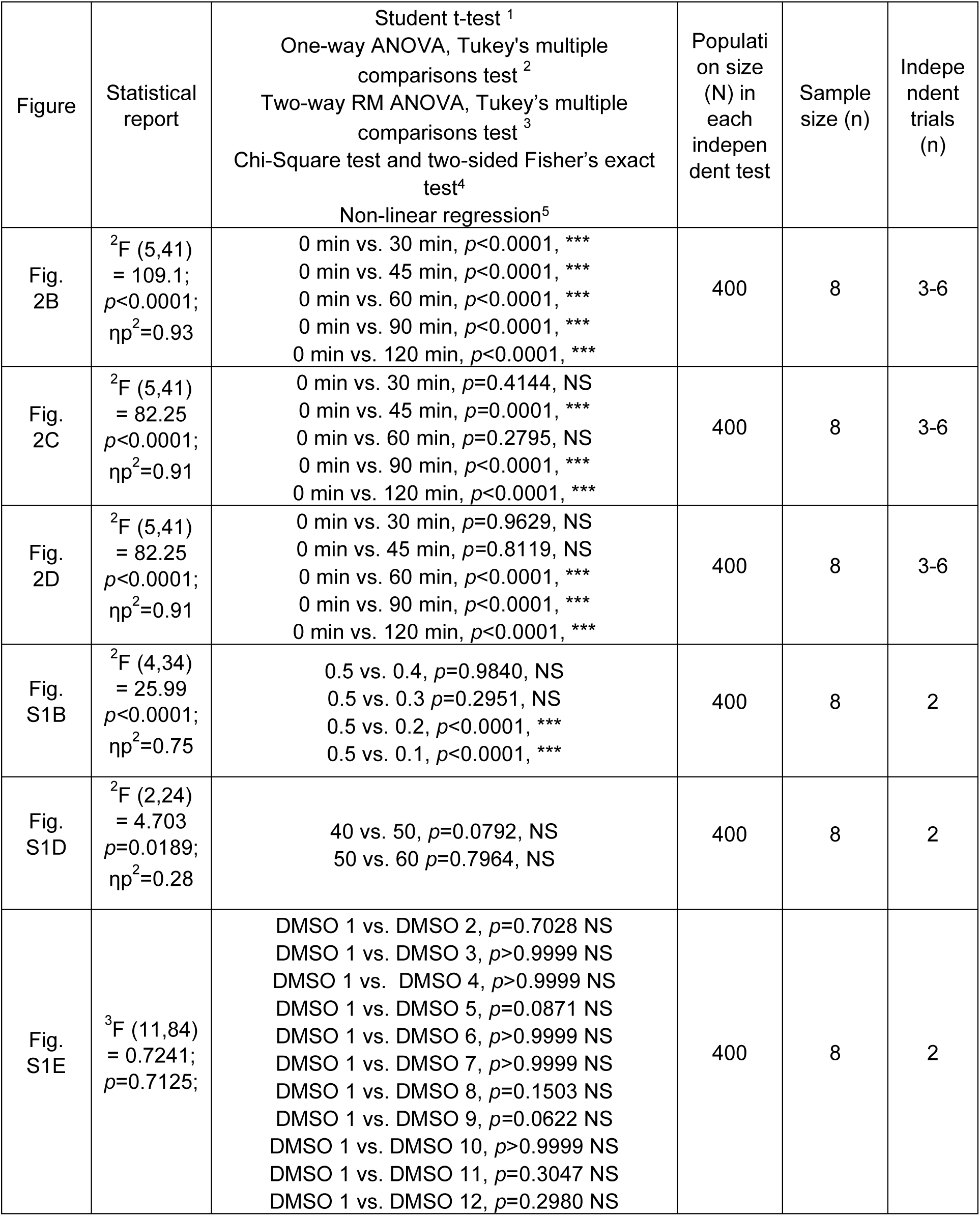
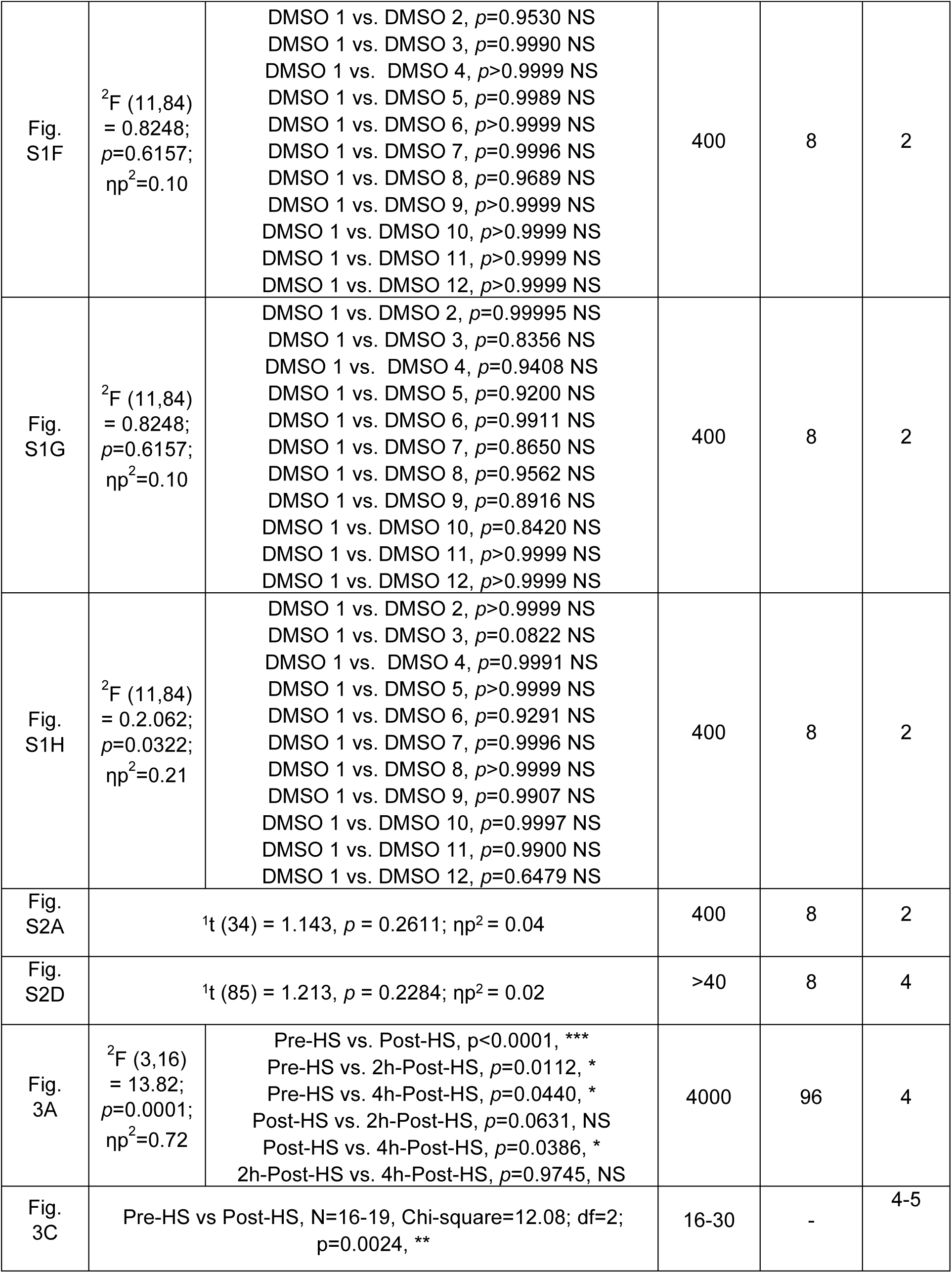
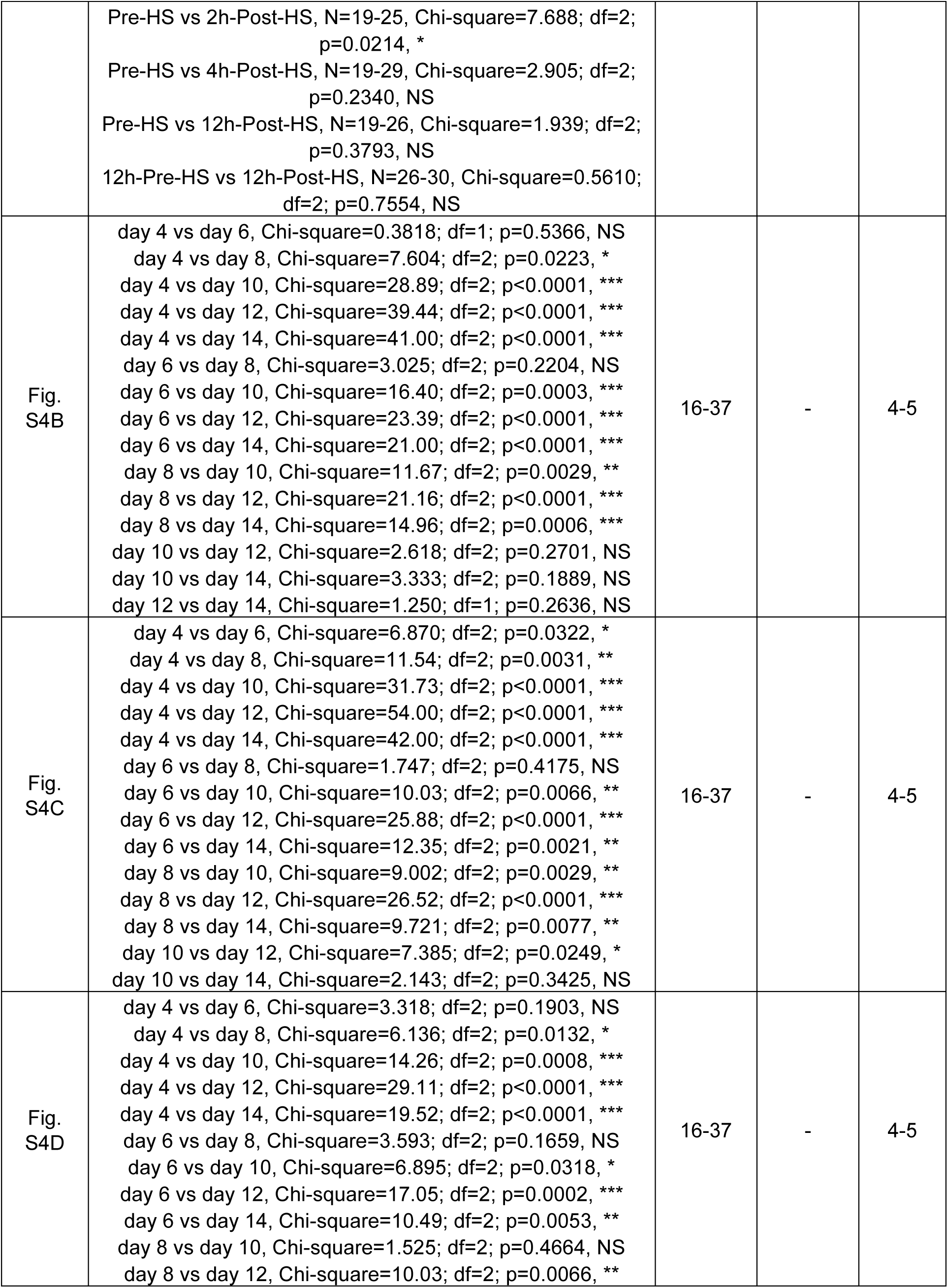
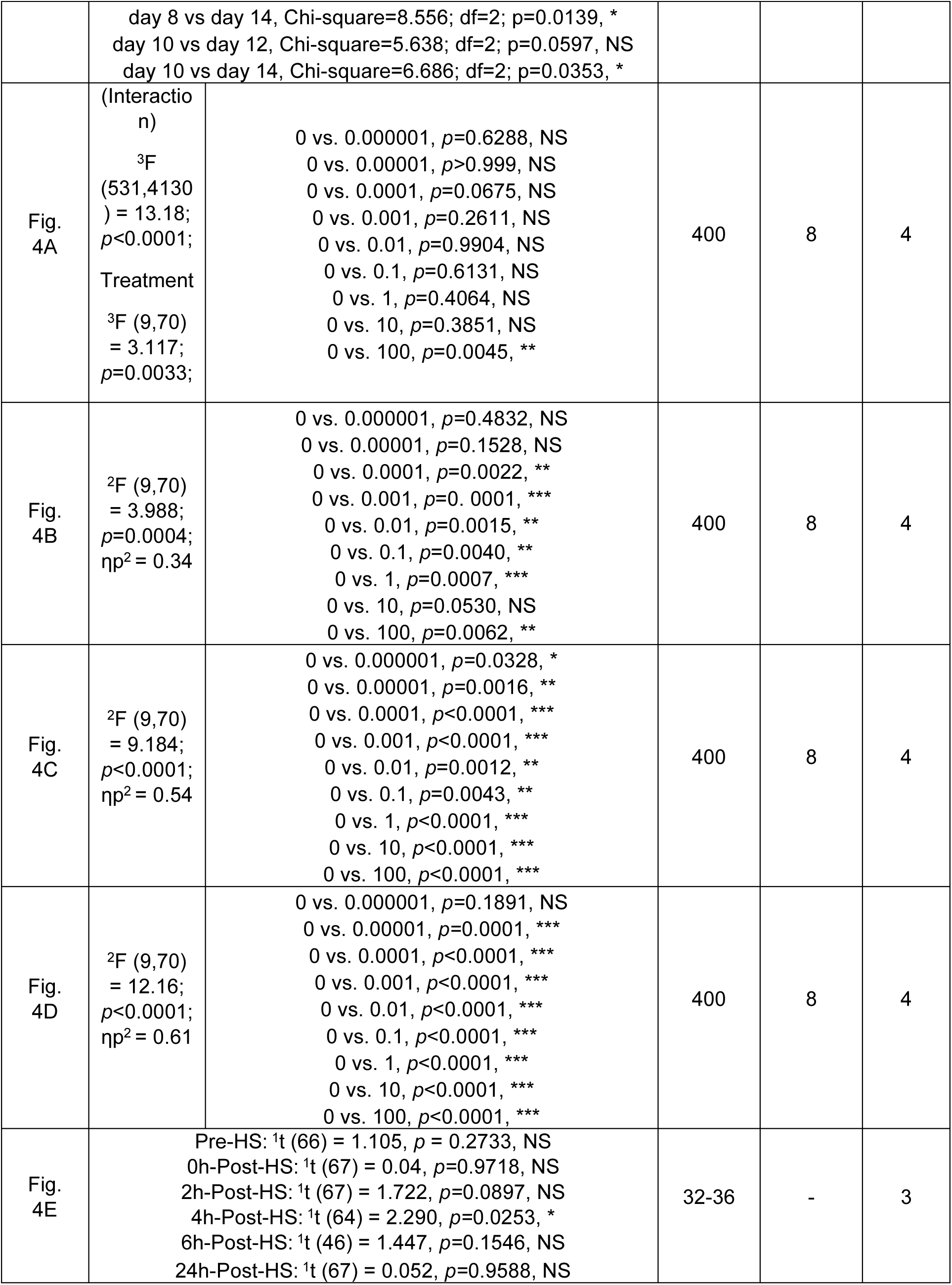
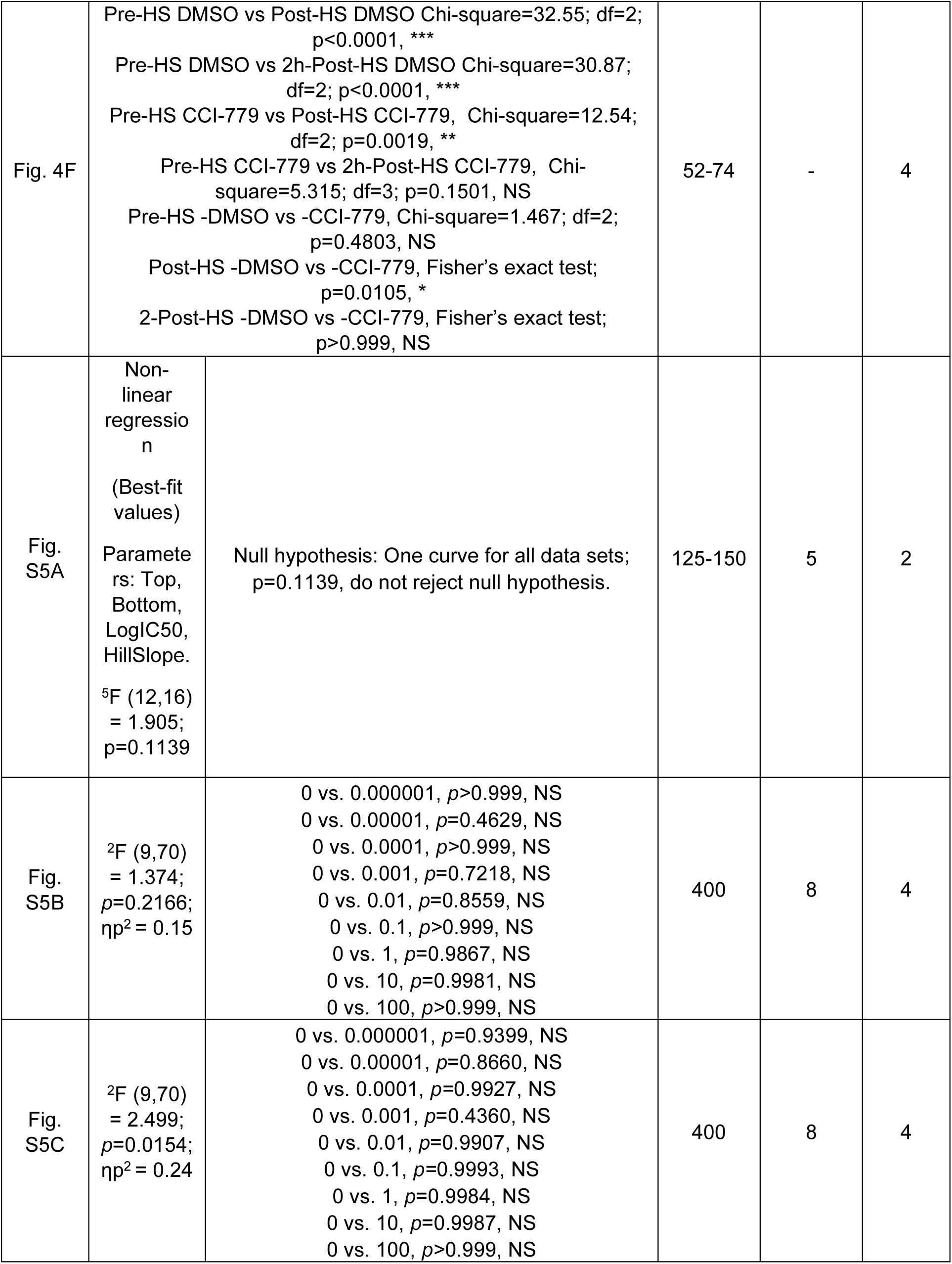
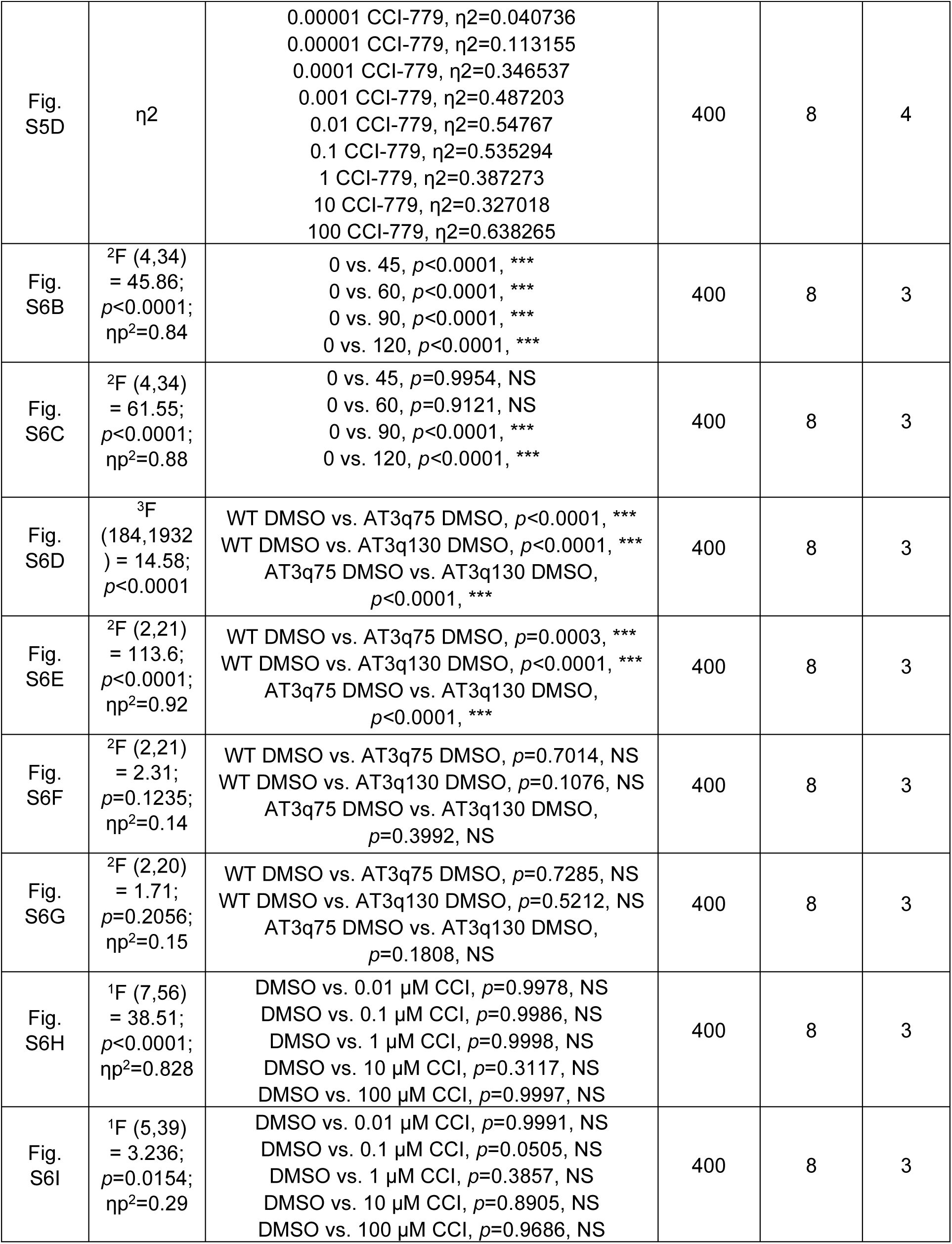
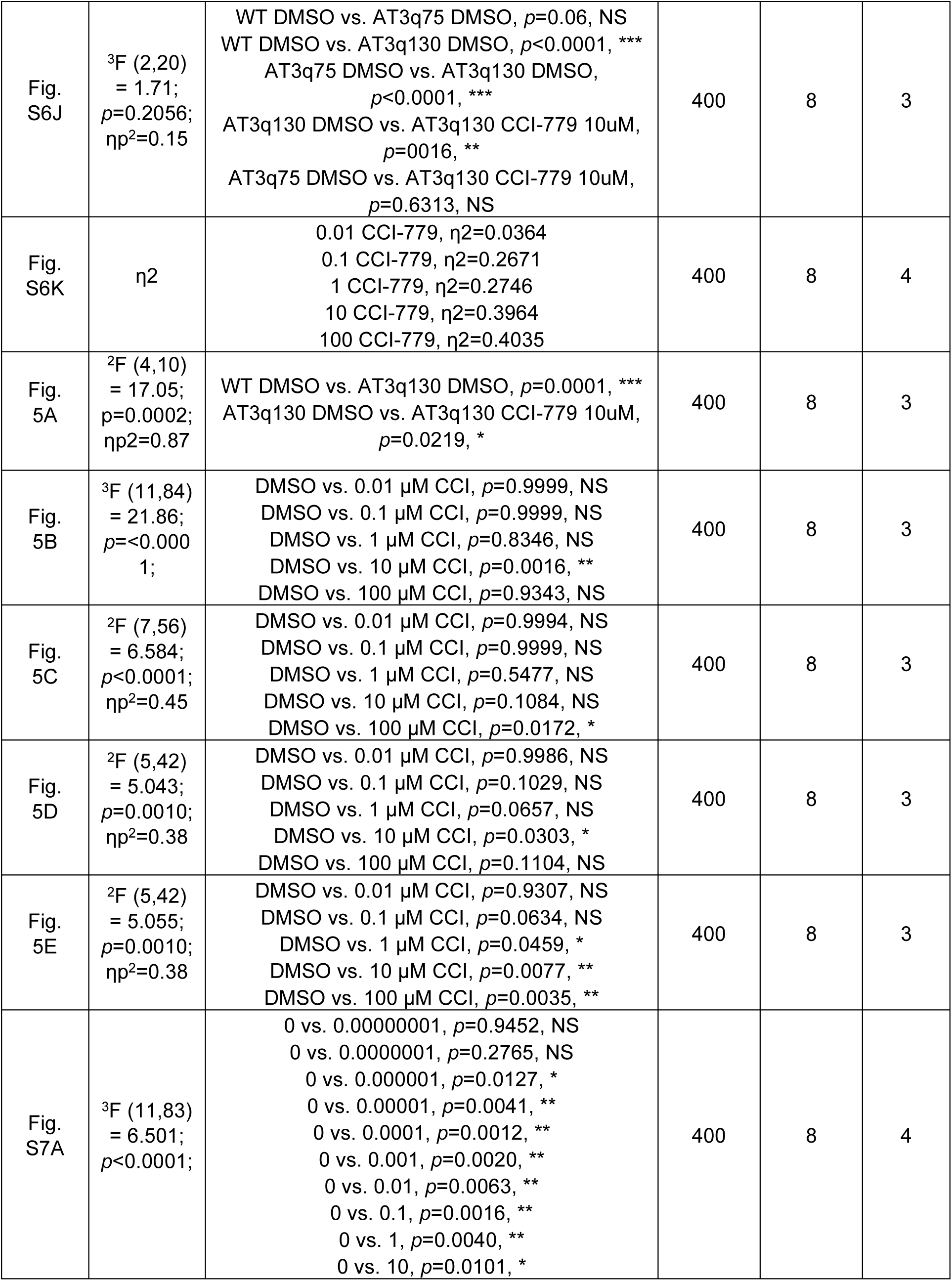
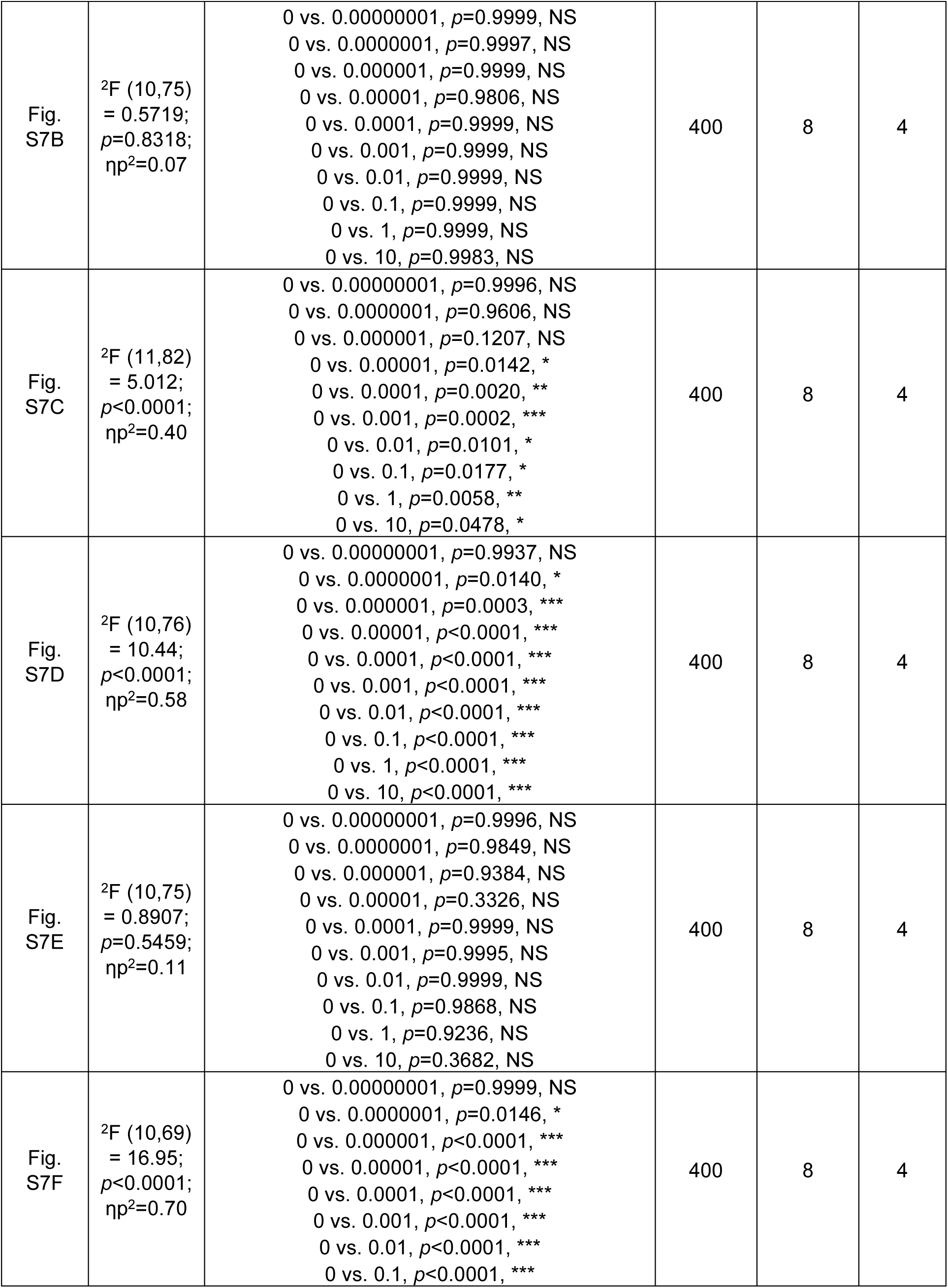
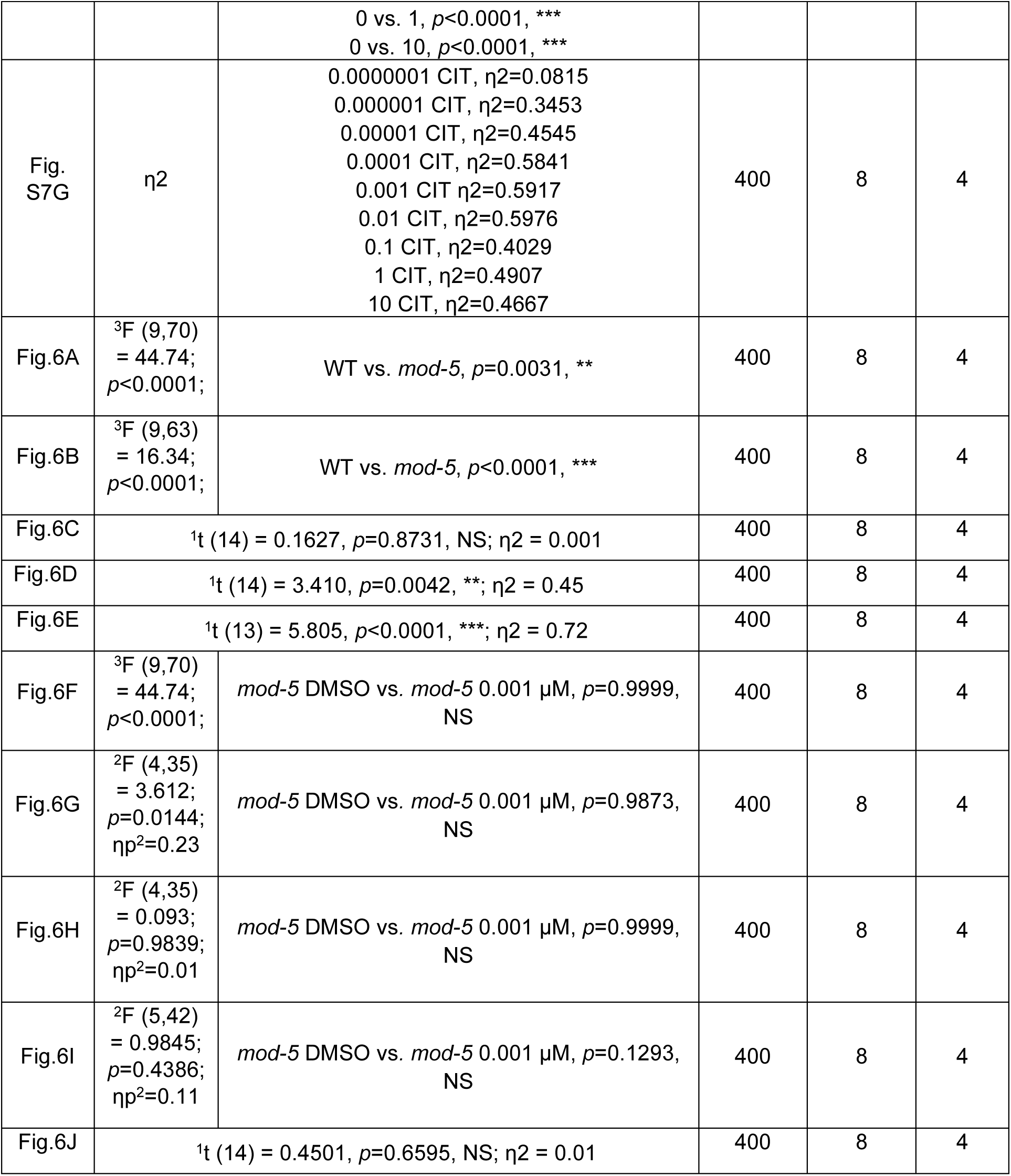
Statistical reports for all experiments.

**Movie S1** (separate file). **Effect of CCI-779 treatment on locomotion recovery in *C. elegans* following HS.** Representative video showing locomotion of WT animals treated with CCI-779 compared to untreated controls at different time points: pre-HS, immediately post-HS, 3 h post-HS, 5 h post-HS, and 15 h post-HS. Animals treated with CCI-779 exhibit accelerated recovery of motor function, with noticeably improved movement as early as 3 hours post-HS, in contrast to controls. These observations mirror the quantitative data shown in Fig. 4A.

**Movie S2** (separate file). **Manual quantification of body bends in *C. elegans* at pre-HS, post-HS, and recovery phases.** The video illustrates representative locomotion behavior of control and CCI-779–treated animals, supporting assay results with an orthogonal method (manual body bend counting, refs. 53, 54). CCI-779–treated animals show increased body bends 2–4 h post-HS compared to controls (see Fig. 4E).

**Movie S3** (separate file). **Effect of citalopram (CIT) treatment on locomotion recovery in *C. elegans* upon HS.** Representative video showing locomotion of WT animals treated with CIT (0.001 µM) compared to untreated controls at different time points post-HS. At 5 h post-HS, CIT-treated animals exhibit markedly improved movement relative to controls, illustrating the enhanced recovery kinetics observed in the assay (see fig. S7).

